# High resolution micro-CT imaging in mice stroke models: from 3D detailed infarct characterization to automatic area segmentation

**DOI:** 10.1101/2022.02.02.478782

**Authors:** Raquel Pinto, Jan Matula, Maria Gomez-Lazaro, Mafalda Sousa, Tomas Zikmund, Jozef Kaiser, João R. Gomes

**Affiliations:** I3S – Instituto de Investigação e Inovação em Saúde, Universidade do Porto, Portugal; Molecular Neurobiology, IBMC-Instituto de Biologia Molecular e Celular, Porto, Portugal; INEB - Instituto de Engenharia Biomédica, Porto, Portugal; Advanced Light Microscopy Unit, I3S – Instituto de Investigação e Inovação em Saúde, Universidade do Porto, Portugal; Central European Institute of Technology, Brno University of Technology, CEITEC-BUT, Czech Republic

**Keywords:** micro-CT, tMCAO, stroke, osmium tetroxide, inorganic iodine, TTC

## Abstract

Characterization of brain infarct lesion in rodent models of stroke is crucial to assess the outcome of stroke therapies and to study the disease pathophysiology. However, so far it has been mostly performed by: 1) histological methods, a time-consuming process that lead to significant flaws and tissue distortion; or 2) via MRI imaging, which is faster, yielding 3D information but at high costs. High-resolution micro-CT imaging became, in the last decade, a simple, fast, and cheaper solution, allowing 3D analysis of samples. Here, we describe that high-resolution micro-CT, either using iodine/O_s_O_4_ as contrast agents, can be successfully applied for fine quantification and localization of lesion size and edema volume in preclinical stroke models. We successfully correlated this new approach with the standard histological method TTC. In transient MCAO mouse stroke model, we were able to identify/quantify large lesioned areas (segmented in core and penumbra), up to degenerated finer striatal myelinated fibers in a transient ischemic attack (TIA) mouse model, at different timepoints post-ischemia, through manual and automatic segmentation approaches (deep learning). Furthermore, 3D reconstructions of the whole brain allow for brain atlas co-registration of the specific affected brain areas. Hence, the presented methodology, through iodine/O_s_O_4_ micro-CT imaging, constitutes a valuable advance in tools for precise and detailed assessment of stroke outcomes in preclinical animal studies.

## INTRODUCTION

Ischemic stroke is the leading cause of mortality and disability and the second leading cause of dementia in western countries ^1^. However, and despite all the pre-clinical findings on the stroke animal models, approaches that aim at reperfusion of the affected brain area are the only treatments available in the last decades ^2^. The lack of translation in stroke pre-clinical studies has recently been appointed as the major factor slowing down the implementation of new therapies ^3^. Rodents are nowadays the preferred animal model in nervous system research, due to their similarity to humans in anatomy, biochemistry and physiology and also low cost of maintenance and processing in a laboratory ^4^. In rodents, stroke is the most frequently modelled by transient occlusion of the middle cerebral artery (tMCAO), where a coated monofilament is introduced into the vasculature to cause occlusion of the mentioned artery for as long as the researcher intends it to ^1, 5^. This means that this model can also be used to study the mechanism and outcomes of different severities of the ischemic lesion. Occlusion of the middle cerebral artery (MCA) for small enough periods causes no neurologic deficit in mice, but the occlusion for longer periods leads to specific neuronal loss, mimicking and modelling the clinical consequences of transient ischemic attacks (TIA) ^6, 7^. Meanwhile, occlusion for longer periods models clinical cases of ischemic stroke both in location and consequences. tMCAO causes the permanent lesion to the striatum (infarct core) and penumbral-characteristics to the surrounding cortex. The penumbra is the brain area that, when affected by stroke, has reduced blood flow while still preserving structural integrity ^8, 9^. Increased attention has been recently given to ischemic penumbra, with this area becoming the current focus of many protective/therapeutic strategies ^10^. Early after stroke onset, the penumbra is the largest part of the infarct volume; and if no reperfusion occurs, it evolves to unrecovered brain infract. Studying the specific localization and temporal expansion of this highly dynamic area (also called peri-infarct-area) while also separating it from the infract core, is of major importance ^8, 9^. Precise localization of the lesion is also becoming increasingly important, due to more attention being given to poststroke recovery therapies, both in preclinical and clinical stroke studies. Since demyelination and axonal breakdown play a critical role in poststroke disability, the detailed characterization of the brain lesion at the fibre level (white matter) is increasingly recognized as an important outcome to measure ^11^. Thus, quantifying the volume of the brain white matter over grey matter, either by quantification of microstructural differences or integrity of myelinated fibres is of major importance to assess poststroke recovery ^12, 13^. Characterization of stroke consequence in rodent models of stroke, as with many other disease-specific animal models, have traditionally relied on histological examination. This methodology can provide high-resolution images from sectioned brains using contrast-enhancing staining, immunohistochemistry, or enzyme histochemistry, and also quick (undetailed) ways to calculate infarct volume ^14^. However, these approaches are highly time-consuming and rely heavily on estimation. An additional obstacle is that brain sectioning may lead to significant damage and distortion of the tissue, as well as the disappearance of part of the sample and loss of 3D context in the brain ^15^. The 3D information becomes of upmost importance when considering that 3D lesion context can be correlated with behavioural assessments in order to give rise to more translatable studies ^16, 17^. Histological examination alone cannot provide that information, only 3D imaging techniques can, especially when co-registered with a neuroanatomical segmented brain Atlas ^18–20^. It is now possible to make 3D non-destructive assessments of brain injury, using both micro-MRI and X-ray micro-computed tomography (micro-CT)^21 22 23^. Micro-MRI demonstrates greater white/grey matter contrast and allows for *in vivo* soft tissue scanning (allowing for longitudinal monitoring of individuals). MRI is currently also the only imaging technique that can distinguish lesion core and penumbra areas in both clinical and pre-clinical research ^24, 25^. MRI imaging, together with classical histopathology, have been used to assess white matter modification in mouse models, while only a few approaches with micro-CT have also been successfully attempted ^26, 27^. However, MRI has poorer spatial resolution, in many cases with anisotropic voxels, which limits the ability to resolve brain fine structures and complicates analysis and interpretation of the 3D data. Furthermore, MRI is more expensive and time consuming than the use of micro-CT imaging. In our work we show that micro-CT is a viable alternative to MRI, mainly due to its high spatial resolution, flexibility and low-cost to the pre-clinical research.

The main disadvantage of micro-CT is the low attenuation of x-rays by soft tissue^28^. This has been overcome using contrast agents stains that enhance soft-tissue x-ray contrast (such as osmium tetroxide (OsO_4_), iodine, phosphomolybdic acid (PMA) and phosphotungstic acid (PTA)) ^16, 29^. Use of these contrast agents has been applied to the brain either by perfusion or immersion, allowing for cerebrovascular or cellular visualization in micro-CT ^30^. The vasculature can be studied using polymerising compounds or agents that penetrate the vessels lumen^31–33 34–36^, meanwhile, neuronal tissue contrast can be achieved through immersion of the whole brain Different contrasts can be achieved because of the different binding affinities of the stains to the different brain cells ^23^. The staining of soft tissues with ionic contrast agents has been used for imaging in the micro-CT of brain ^37^ and whole-body^38^ of zebrafish, as well as, rodent embryos, brains and other organs ^28, 29, 38^. Some have even gone further, staining whole mice brains to identify/quantify electrode lesions ^39^, tumours^40^, malformations ^41^, disease pathology in Alzheimer’s ^42^ and even single neurons^43 44^. Nevertheless, the use of these stains for the visualization and quantification of cerebrovascular alterations ^45^ and ischemic lesion ^16, 46–48^ in whole mouse brains have rarely been described and never with great tissue/lesion contrast.

While sample preparation and micro-CT measurement itself constitute major steps in the micro-CT analysis, it is only the beginning of the whole pipeline. Image segmentation is the next crucial step, where analysed structures are defined by assigning a class to each pixel/voxel of an image or volumetric data. This is a particularly difficult task in the segmentation of specific brain structures, due to the low contrast offered by different types of tissue and anatomical structures. Manual segmentation, where a skilled operator delineates the structures of interest by hand, is thus still popular in the segmentation of micro-CT images of the brain, due to its accuracy and low requirements ^16, 42^. However, it’s major disadvantage is the high time-cost, which poses a significant bottleneck in the whole analysis pipeline. Manual segmentation also introduces operator bias, which can affect the accuracy of subsequent analysis. These issues become even more prominent in large-scale studies with many brain samples analysed. The focus is thus shifting towards the development of semi-automatic and fully automatic tools, which make the image segmentation significantly faster. Deep learning and specifically convolutional neural networks (CNNs) have recently seen an enormous increase in popularity for their application in difficult biological micro-CT image segmentation tasks ^49^. CNN has been recently used for segmentation of: ant brain ^50^, murine mineralised cartilage/bone ^51^ and rabbit calcified knee cartilages ^52^ in micro-CT images. While this segmentation provides us with morphological information about the object of interest, it does not tell us anything about its function in the context of brain anatomy. Coupling micro-CT spatial information with an anatomical atlas can provide accurate location of stroke-affected areas in the context of the whole brain and area functions^53^.

Thus, in this study we show how the use of high-resolution micro-CT can improve the assessment of stroke outcomes in brain rodent models. We describe: 1) a simple and effective *ex vivo* staining method for the brain using two contrasting agents, Iodine and Osmium tetroxide; both stains originate images with a high-quality, resolution, and neuroanatomical detail; 2) a novel methodology that allows precise quantification of 3D brain lesion, from brain edema to total infarct area, with core/penumbra segmentation; 3) the successful validation of this new approach as compared to classical histological evaluation, TTC; 4) a semi-automatic method to evaluate white matter/grey matter changes in integrity of myelinated fibres; 5) a novel automatic, deep learning-based, method for segmentation of the lesion and penumbra in the micro-CT images; and 6) co-registration of 3D brain images with mouse atlas, allowing specific brain areas identification.

This way, the complete methodology here described, from the used contrasting agents to the stroke analysis methods, highlight the advantages of micro-CT imaging as a relevant tool for the assessment of preclinical stroke studies.

## RESULTS

### High resolution micro-CT imaging allows brain neuroanatomical areas distinct identification

The complete methodology described in this work was developed from the need to finely characterize and quantify ischemic alterations in mice brains, without the requirement of laborious histological techniques or costly MRI apparatus. As such, whole mice brains were subjected to different staining methodologies for soft tissues, previously described in the literature with limited success (low resolution and detail): osmium tetroxide, inorganic iodine, iohexol (Omnipaque ®), PMA and PTA. X-ray voltage and output current were fine-tuned to optimize volumetric reconstructions for each stain at isotropic resolutions, in order to allow histopathological interpretations (see materials and methods). For whole mouse brain *ex vivo* imaging, the brains were horizontally mounted on the micro-CT apparatus and complete brain imaged. Each brain was reconstructed to obtain a single 3D volume (**Fig.1A**). For high resolution micro-CT imaging, image dataset acquisitions took from 3 to 5 hours, which is much faster than the determination by histological methods, that might take days. In this regard, for large-scale stroke studies with focus on the quantification of the lesion of large number of mice brains our approach facilitates high-throughput analysis.

**Fig. 1.**
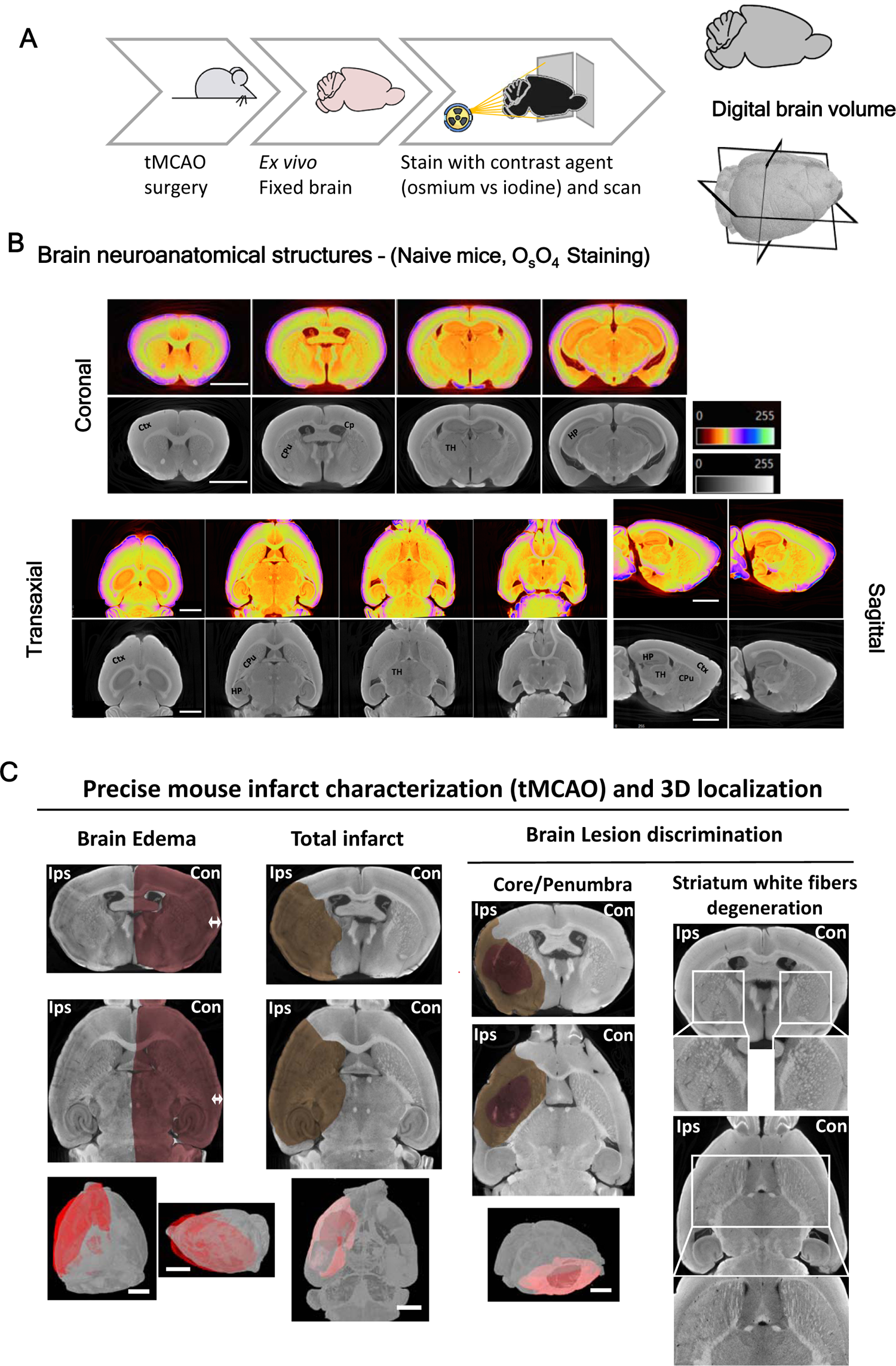
High resolution micro-CT imaging allows brain neuroanatomical areas identification in physiological/pathological conditions (mouse stroke models). (A) Overview of the technical steps required to prepare whole mice brains for micro-CT imaging. (B) Representative images in different projections of the brain (control, naive) showing contrast enhancement using osmium tetroxide stain to reveal brain neuroanatomical structures, and clear white/grey matter distinction. Regions such as cortex (Ctx), hippocampus sub-areas (HP, CA1-3, DG), choroid plexus (Cp) and striatum fibers in Caudate putamen (CPu) and thalamus (TH) among others, can be identified. (C) Representative 3D images of the application of osmium staining to evaluate features induced by mouse stroke model (tMCAO) and TIA models, including: brain edema (arrow indicates the brain swelling, comparison between contra/ipsilateral sides), total infarct area (including core (highlighted darker area)/penumbra ((highlighted less dark area) discrimination), and striatum white fiber degeneration. Scale bar: 2mm.

From all the compounds tested PTA and PMA were not able to stain the whole brain and seemed to not penetrate completely in the tissue (**Supplementary Fig. 1B,C**). In the case of iohexol, despite its ability to penetrate the whole sample, it conferred poor contrast of the different brain structures (**Supplementary Fig. 1A**). Meanwhile, brains stained with osmium tetroxide (**Fig. 1B**) and the inorganic iodine (**Fig. 3**) showed high brain structural contrast in scans at resolutions of 3.5-4 µm voxel, allowing for a microscopic reconstruction of the whole-brain 3D volume (**Fig. 1A,B** - coronal, sagittal and transaxial views).

Importantly, the two successful stains (osmium tetroxide and iodine) produced complementary micro-CT contrast: since inorganic iodine is soluble in water, it enhances contrast mainly in structures of the grey matter and myelin; whereas osmium is lipophilic, providing increased contrast to cell membranes and other lipid-rich structures, and creating a rather homogeneous contrast to all cell types^22 38 26^. Although iodine stain resulted in higher grey/white matter contrast, both compounds allow a clear distinction of several brain structures such as cortex (Ctx), striatum (CPu), hypothalamus, thalamus (TH) and hippocampus (HP) (**Fig. 1B** and **Supplementary Fig. 3**). Iodine has also an important advantage regarding toxicity over osmium (**Table 1**).

**Table 1.** Comparisons of the different characteristics of osmium tetroxide vs inorganic iodine staining for whole-brain contrast enhancement.

|  | Osmium tetroxide | Inorganic iodine |
| --- | --- | --- |
| Binds lipophilic structures | ++ | - |
| Binds hydrophilic structures | - | ++ |
| Toxic | ++ | - |
| Expensive | ++ | - |
| Allows further histological processing | - | ++ |
| Penetration speed | + | ++ |
| Brain shrinkage | - | ++ |
| Edema extent quantification | ++ | ++ |
| Lesion (core/penumbra) volume quantification | ++ | ++ |
| Striatum white matter degeneration quantification | + | ++ |
| Lesion (core/penumbra) volume quantification | + | ++ |

**Stroke-related lesions can be observed on mouse brains using high-resolution micro-CT Mouse** brains subjected to ischemic insult were processed for tissue staining with either osmium tetroxide or iodine. After micro-CT image acquisition the datasets were reconstructed from the x-ray projections. Despite the differences in image contrast, both osmium tetroxide and iodine stained brains revealed clearly recognizable effects from the ischemic injury (**Fig. 1C, 2A – osmium and Supplementary Fig. 3, iodine**), which includes brain edema and degeneration in the stroke area. At the site of lesion, differentiation between core and the penumbral area was evident at 24-hours post-reperfusion (**Fig.2A-C, Suppl. Video 1)**. Additionally, striatum fiber degeneration in the TIA model was perceptible (**Fig. 3A-C**). Advantageously, iodine-stained brains could be used for further histological processing since, after the staining process, the tissue retains the structural components for the recognition by antibodies used in immunofluorescence protocols (**Supplementary Fig. 2B**). This correlated multimodal image acquisition workflow contributes to the lower number of animals needed for experimentation and allows to obtain morphological information and, when imaged at higher resolution with wide-field or confocal microscopy, for the acquisition of functional information. However, due to steps of dehydrations performed in the protocol for iodine staining, brains suffered some shrinkage. The final brain volume was calculated to be a third of the initial volume (**Supplementary Fig. 2A**). Opposite, although brain shrinkage does not occur with the brains stained with osmium tetroxide, the staining process leads to tissue mineralization^54 37 23^, further preventing its use for immunofluorescence. However, staining with toluidine blue can still be successfully performed (results not shown), which allows for DNA and proteoglycan staining ^55^. In **Table 1** the advantages/disadvantages of the two contrasting agents (osmium tetroxide and inorganic iodine) are described for whole-brain contrast enhancement.

**Fig. 2.**
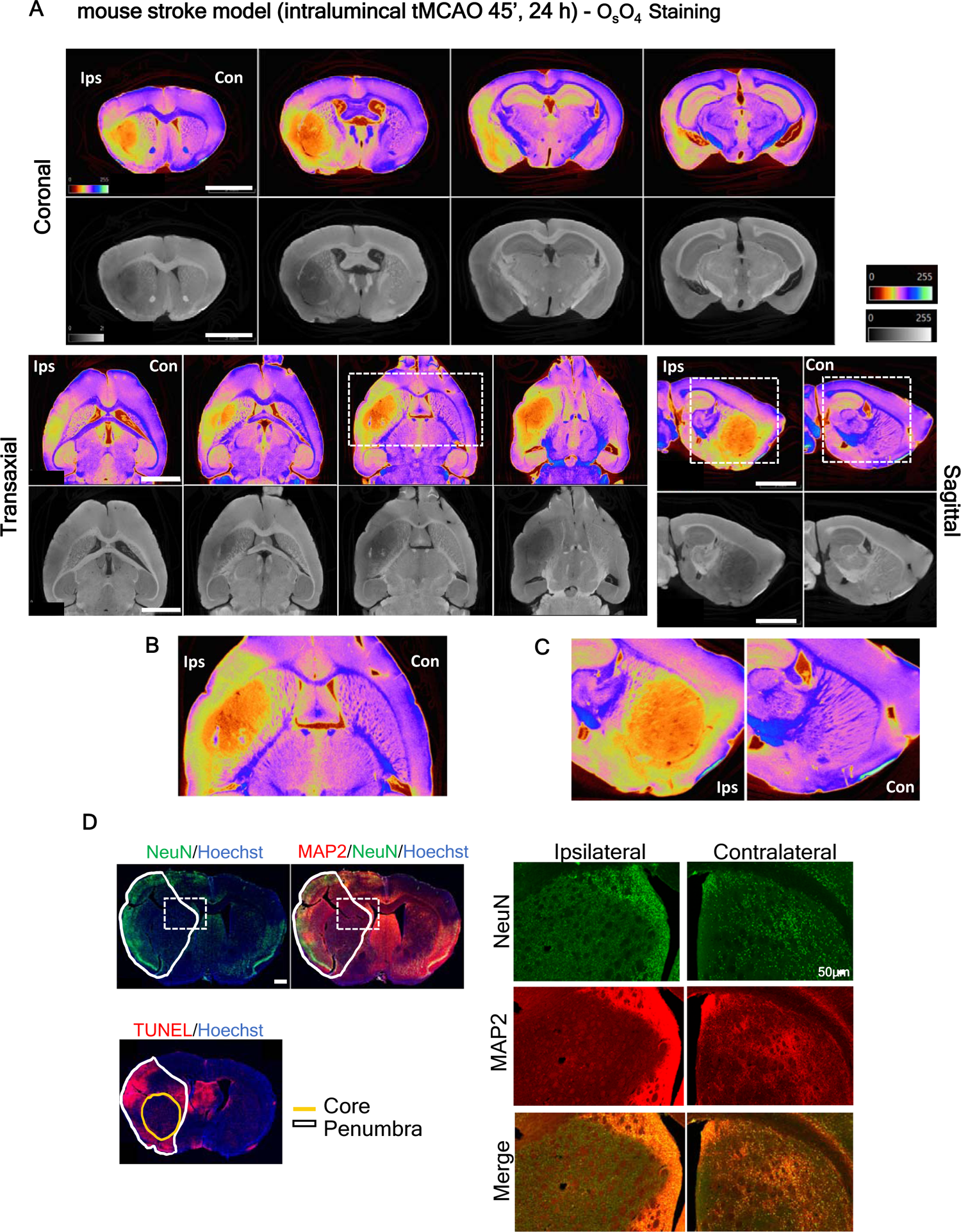
High resolution micro-CT imaging allows clear visualization of mouse stroke model (tMCAO) lesions. **(A)** Virtual slices (∼4µm voxels) at arbitrary orientations from micro-CT scan of a mouse stroke brain immerged in osmium tetroxide showing total lesion as well as core (orange/darker) and penumbra ((yellow/grey around core) differentiation. Below panel shows lesion detail in the **(B)** transaxial and **(C)** sagittal pane. Scale bar: 2 mm. **(D)** Representative brain slices immunolabeled with specific neuronal markers (NeuN and MAP2), as well as with a marker for apoptosis -TUNEL, confirming that lesion demarked area in micro-CT images corresponds to the histologically defined lesion. Scale bar: 500 µm. Right side panel shows a lesion in detail in the coronal plane, Scale bar: 50 µm.

**Fig. 3.**
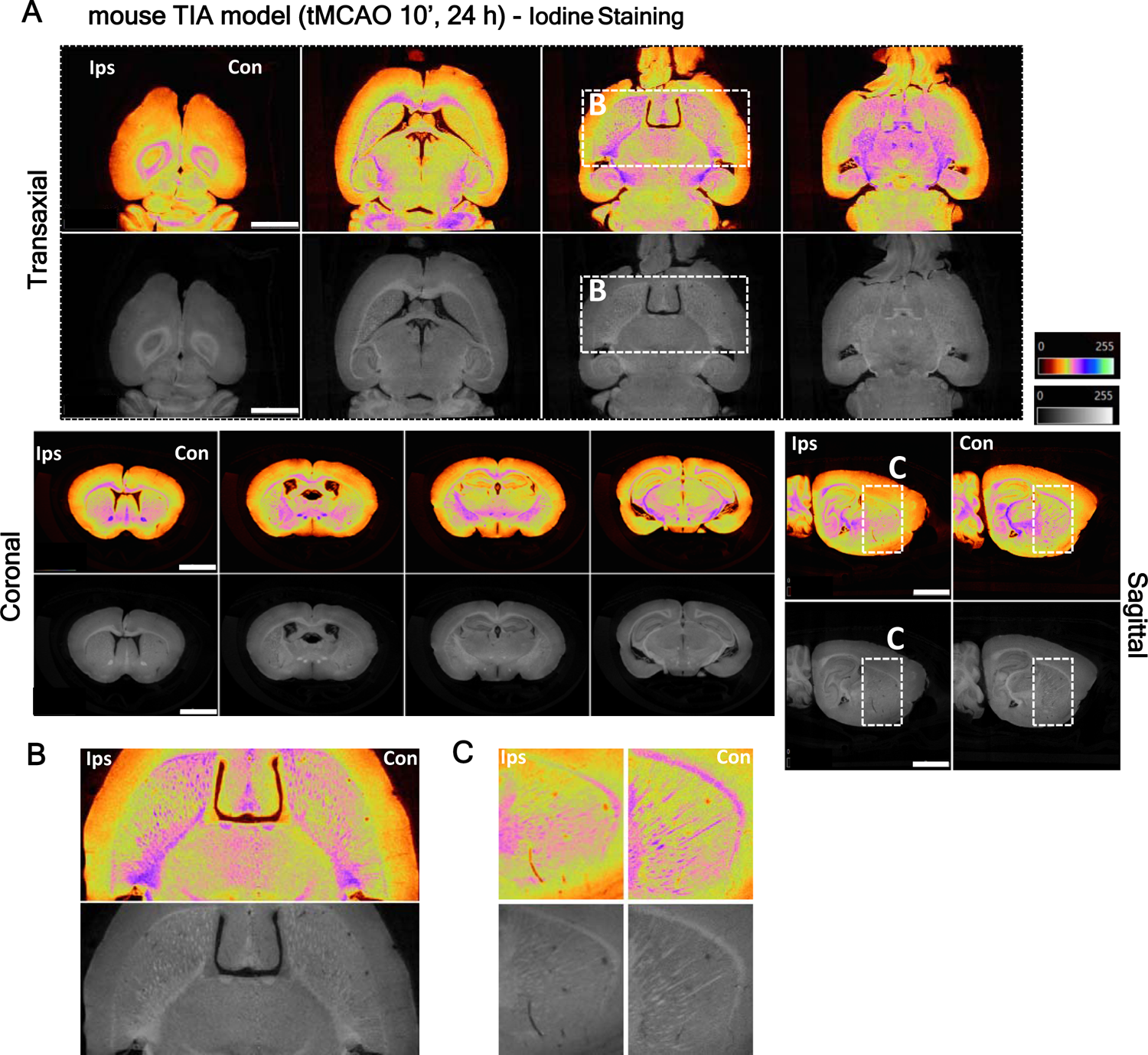
High resolution micro-CT imaging can identify striatum white fiber degeneration in transient ischemic attach (TIA) mouse model lesions. (A) Virtual slices (∼4µm voxels) at arbitrary orientations from micro-CT scan of a TIA brain immerged in inorganic iodine, showing degeneration of striatum white fiber matter, after 10min MCA occlusion. (B,C) Below panel shows lesion detail in the transaxial and sagittal plane. Scale bar: 2mm.

### Characterization of mice brain infract lesions

Successful stroke therapy assessment requires the detection of subtle differences in infarct volume. By optimizing contrast-enhanced micro-CT imaging (O_s_O_4_ or iodine), we demonstrated that this methodology is suitable for defining ischemic lesion boundaries, including discrimination of ischemic core from penumbra at 24h following reperfusion, and visualization of brain edema (**Fig. 2A and supplementary Fig. 3**), as well as following brain lesion over-time, from 3h to 72h (**Supplemental Fig. 5**), till 7 days (tested, data not shown). With the described methodology it was possible to observe a time dependent increase in infarct volume in the cortex/striatum, as well as an increase in ipsilateral hemispheric volume due to edema (from 3 to 24 h after reperfusion) (**Supplementary Fig. 5**). The area of the lesion, area with lowest x-ray contrast, was calculated with the 3D image datasets obtained with the micro-CT. These values were compared with the information obtained by using the standard Triphenyl tetrazolium chloride (TTC) staining (**Fig. 8**), together with the information obtained by immunolabeling for NeuN (neuronal marker), MAP2 (neurites marker) and TUNEL (late apoptosis/necrosis marker) (**Fig. 2D**). From our analysis, the area of the lesion calculated from the virtual slices was in close agreement with the area of the lesion quantified by histological sectioning, confirming that the described demarcation corresponds to the ischemic infarction area (**Fig. 2D and Fig. 8C,D**). Additionally, core/penumbra discrimination can be achieved in early timepoints (24h) ((**Fig.2B and Supplementary Fig. 5)**. Moreover, micro-CT imaging allowed the detection of small lesions in myelinated fibres after the mouse TIA model. The contrast conferred by the stains (Osmium, **Supplementary Fig.4, Suppl. Video 2; Iodine Fig. 3**) enabled the visualization of white matter loss in the affected hemisphere’s striatum, when compared to the contralateral hemisphere after 24h reperfusion. White matter fiber tracts loss was uniform throughout the striatum, and not localized to regions of ischemic injury, since this model does not present a visible infarct area (**Fig. 3 and Supplementary Fig. 4**). Thus, both osmium and iodine ex-vivo stained brain’s show clear stroke consequence demarcation that parallel histopathological-defined ischemia.

### High resolution micro-CT allows fine manual quantification of mouse brain lesion/edema

The micro-CT proprietary software from Bruker®, CTAn, was used to manually delineate the regions of interest of both hemispheres and the ischemic lesions, including core when present, in the virtual slices (**Fig. 5**). The selected regions of interest were further used to calculate edema extent (% contralateral hemisphere) (**Fig. 4A-C**) and corrected lesion volume (% of total brain volume) (**Fig. 4D-F**) (see materials and methods section). Moreover, this approach offers several advantages over classical histological methods since: the area occupied by the ventricles can be included/excluded in the calculations depending on edema extent; brain midline, which shifts due to edema, can be adjusted allowing fine brain area selection. Osmium tetroxide-stained brains subjected to the 45-minutes occlusion (stroke model) displayed an edema extent that lead to an increment of the size of the ipsilateral hemisphere, 10.6% ± 3 %, when compared to the contralateral hemisphere, and exhibited lesions that occupied 13.7% ± 1% of the total brain volume (penumbra: 12.07% ±0.7 % and core: 1.6% ± 0.5 %), while iodine-stained brains of the same stroke condition displayed an edema extent of 5.1% ± 2 % and total lesion of 14.5% ± 2 % (penumbra: 13.5%± 1%, core: 0.9% ± 0.8%) (**Fig. 4C,F**). In this way, both staining methods demonstrated to be useful for the quantification of brain edema and stroke areas.

**Fig. 4.**
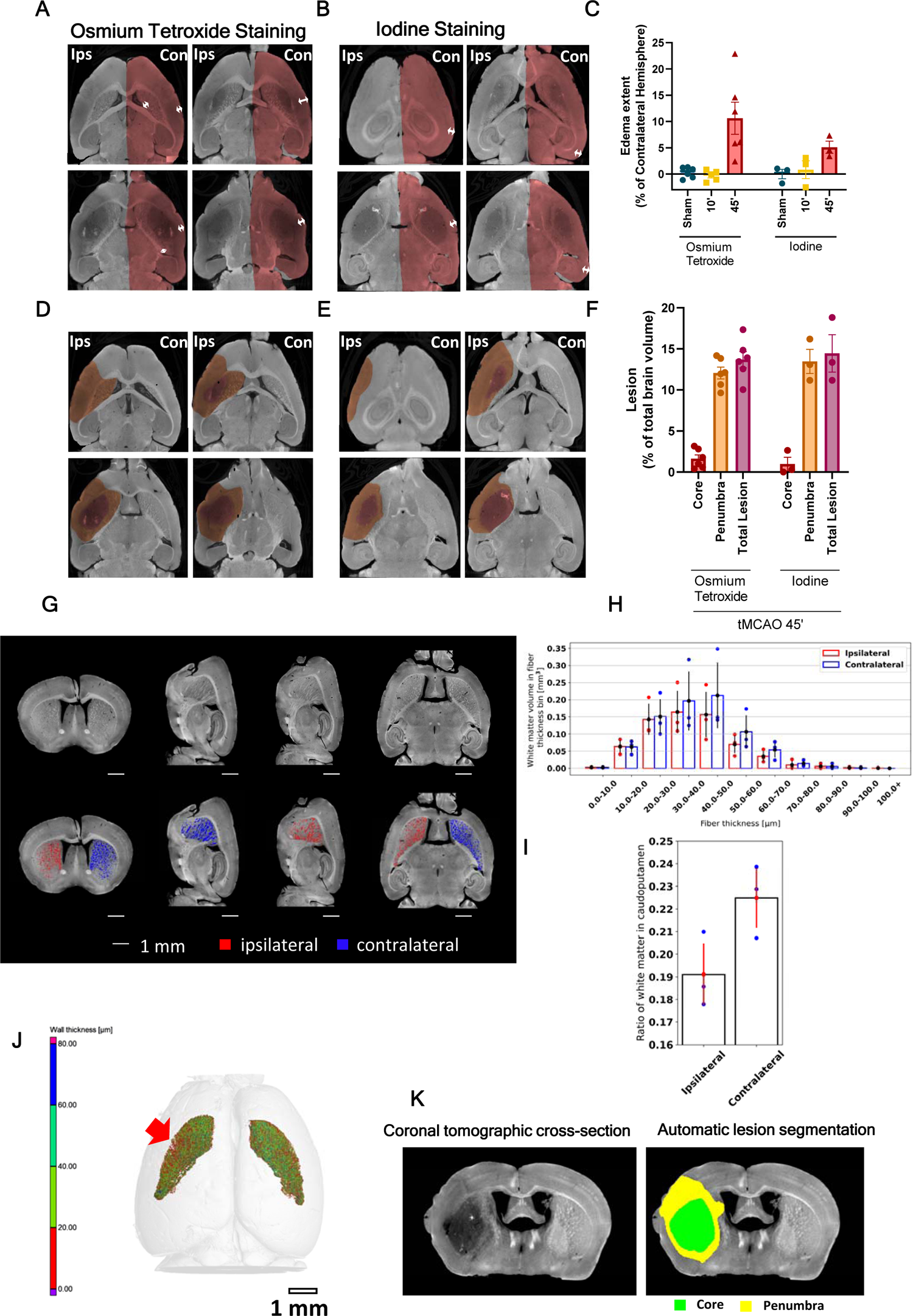
Micro-CT imaging allows fine quantification and characterization of tMCAO (stroke and TIA) lesions in mice models. 2D slices in the transaxial plane showing overlayed (A,B) edema (arrows indicate the brain swelling differences, between contra/ipsilateral sides) and (D,E) lesion (highlighted less dark area) /core (highlighted darker area) annotation for each staining used: (A,D) osmium tetroxide and (B,E) inorganic iodine. (C) tMCAO brain’s edema extent, as quantified using the manually obtained volumes for each brain hemisphere in the micro-CT images. (Osmium tetroxide: sham, n=6; 10-minute, n=5; 45-minute, n=7. Inorganic iodine: sham, n=3; 10-min, n=3; 45-min, n=3). (F) Edema-corrected lesion volume (with discrimination between core and penumbra), as quantified using the edema extent and the total lesion/core volumes, manually segmented (Osmium tetroxide, tMCAO 45-minute, n=6. Inorganic iodine, tMCAO 45-minute, n=3). (G) Representative images of iodine-stained brains from TIA mouse model showing evident white matter contrast at arbitrary orientation and white matter degeneration in caudate putamen ipsilateral hemisphere. (H) Automatic quantification of white matter volume difference between hemispheres] in different individual fiber thicknesses (n=3, A-C). (I) Ratio of caudate -putamen volume occupied by white matter in each hemisphere (ipsilateral vs. contralateral hemisphere), n = 3. (J) Heat map of striatum white matter wall thickness, as quantified and segmented by the semi-automatic method, red arrow shows an area of significant decrease in fiber thickness in the ipsilateral hemisphere (of TIA model). (K) Representative images of iodine-stained brain of mouse stroke model (tMCAO 45’) and after automatic lesion segmentation.

**Fig. 5.**
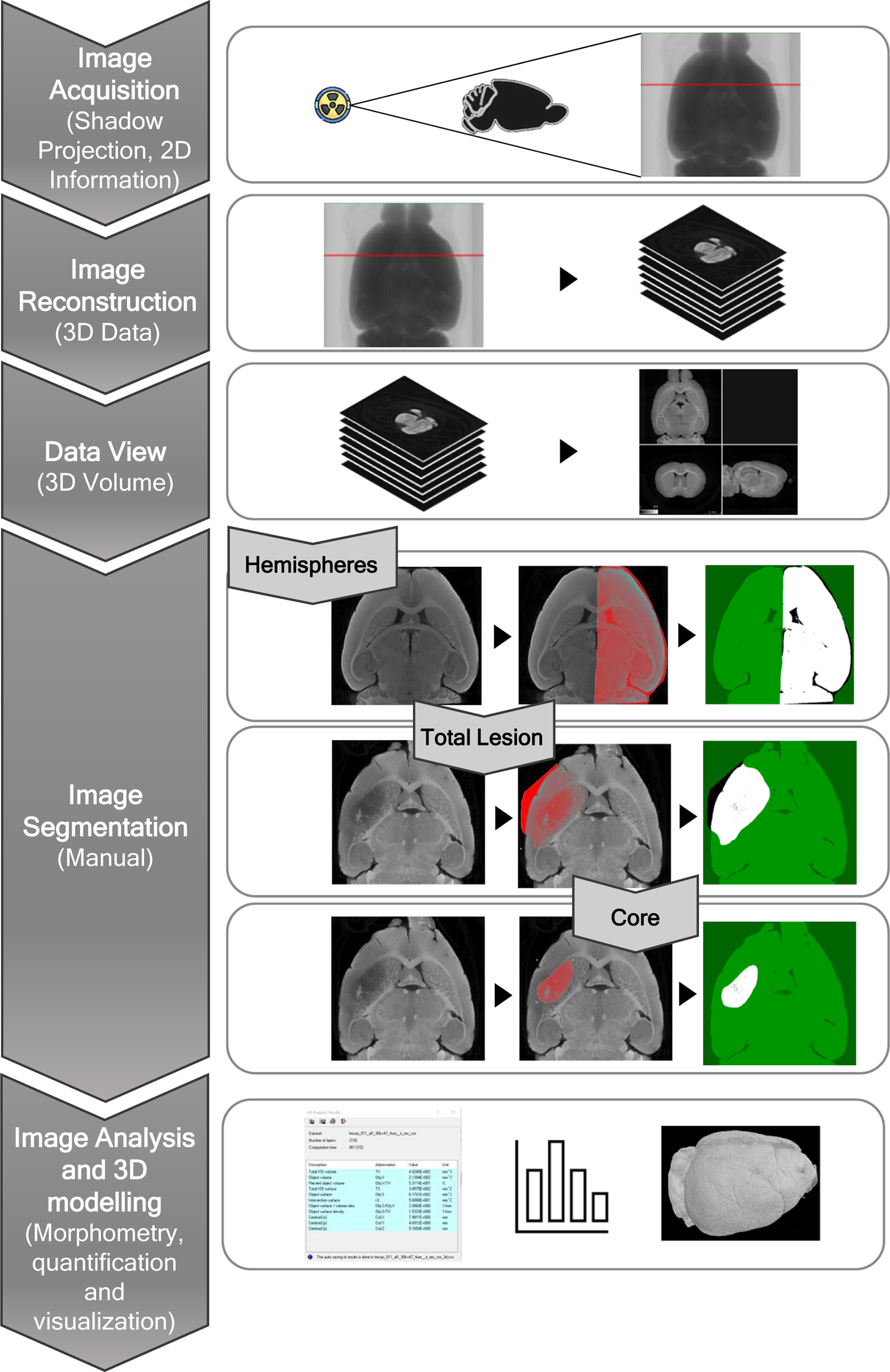
Manual quantification workflow of the different ischemic parameters. Representation of manual quantification procedure of mice brains subjected to stroke models and stained with a contrast agent. Images were reconstructed for 3D rendering using the NRecon software from Bruker and visualized in the DataViewer Bruker software. Analysis was performed in the CTAn software from Bruker, in which hemispheres, total lesion and core were manually delineated on the transaxial plane, and volume occupied by the respective regions of interests calculated.

### Deep learning enables fully automatic lesion segmentation in the micro-CT images

To improve the reproducibility, high throughput quantification and reduce operator bias, we designed an automatic segmentation process to quantify micro-CT lesions. This is a difficult task due to the low contrast between the lesion and healthy tissue. However, iodine-stained brains from the stroke model, which display higher signal to noise ratio, allowed for the application of a fully automatic deep learning-based segmentation and quantification method to define ischemic lesion, with both core and penumbra (**Fig. 4J/K, Fig. 6B**). The method is based on training a convolutional neural network to transform a lesioned brain hemisphere to an approximation of a healthy brain hemisphere by randomly simulating the lesion in the available micro-CT cross-sections of healthy brain hemispheres. By subtraction of the original and transformed image and subsequent thresholding of this difference image, the segmentation masks of the whole lesion and core are obtained. The results of this automatic quantification in iodine-stained brains of the tMCAO model are shown in **Table 2**. The total lesion quantified by this method was 20.33 ± 4.48 mm^3^ while the core lesion was 2.36 ± 1.56 mm^3^, matching the manually obtained volume measures of 21.63 ± 4.75 mm^3^ for the total lesion volume and 3.10 ± 1.39 mm^3^ for the core volume. Micro-CT in combination with deep learning can thus be utilized for fast evaluation of the lesion morphology in 3-D.

**Fig. 6.**
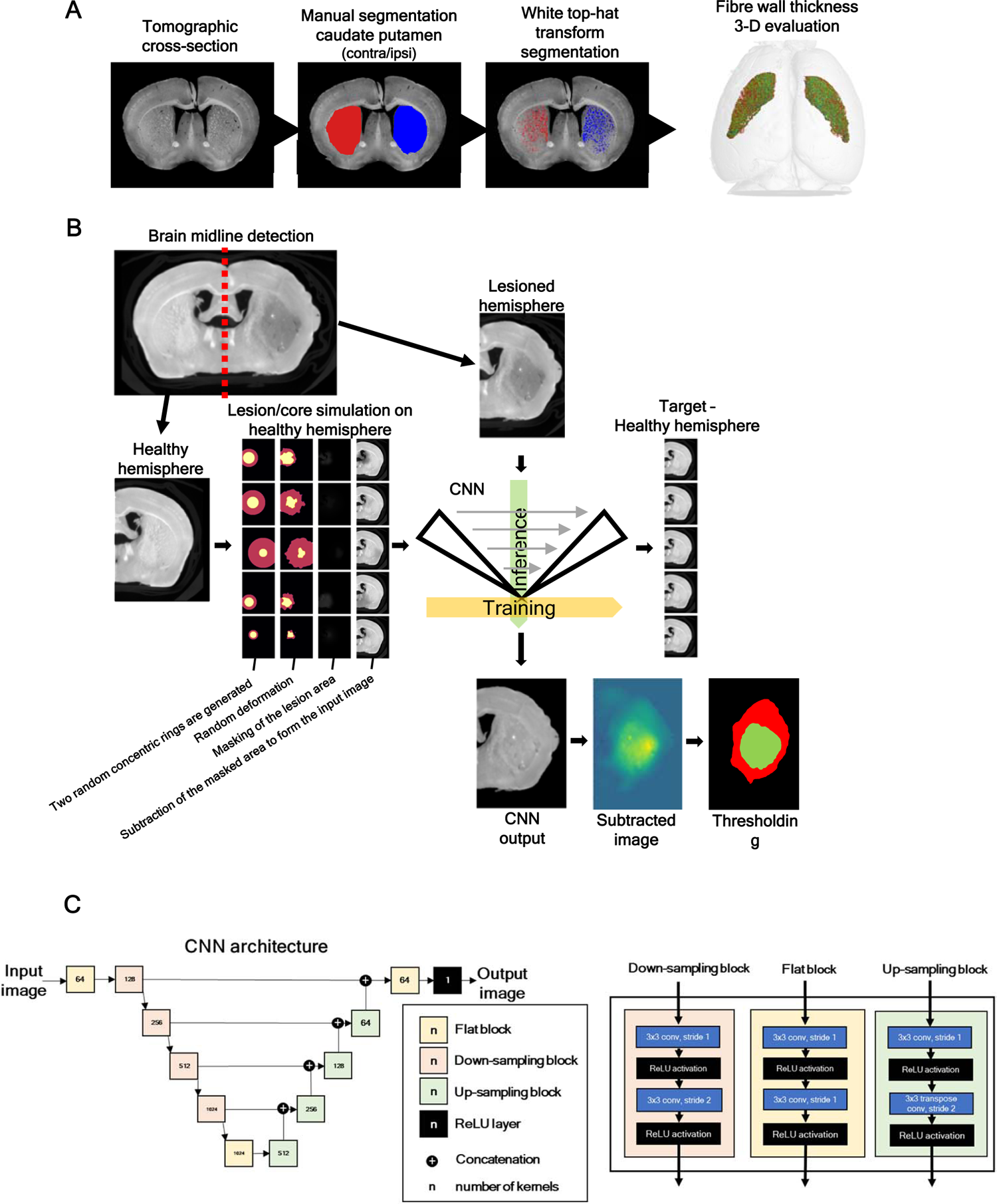
Proposed methods for semi-automatic evaluation of the white fiber degradation in TIA model. (A) and automatic infarct lesion segmentation, in stroke model (B). In (A) combination of manual segmentation of caudate putamen and segmentation of white matter fibers via white top-hat transform was utilized. The thickness of the fibers was then evaluated in the software VG Studio MAX 3.4 (Volume Graphics GmbH, Heidelberg, Germany). In (B) an automatic deep learning-based segmentation workflow is proposed for the segmentation of the lesion penumbra and core. (C) The CNN utilized for the automatic lesion detection.

**Table 2.** Results of automatic quantification of iodine-stained tMCAO brains based on deep-learning-based segmentation vs manual segmentation.

|  | Automatically segmented lesion volume (mm<sup>3</sup>) |  | Manually segmented lesion volume (mm<sup>3</sup>) |  |
| --- | --- | --- | --- | --- |
|  | Total Lesion | Core | Total Lesion | Core |
| Sample 1 | 16.8906 | 3.7642 | 20.2020 | 5.0047 |
| Sample 2 | 17.4301 | 3.1330 | 16.6580 | 2.5465 |
| Sample 3 | 26.6595 | 0.1852 | 28.0330 | 1.7491 |

**Table 3:**
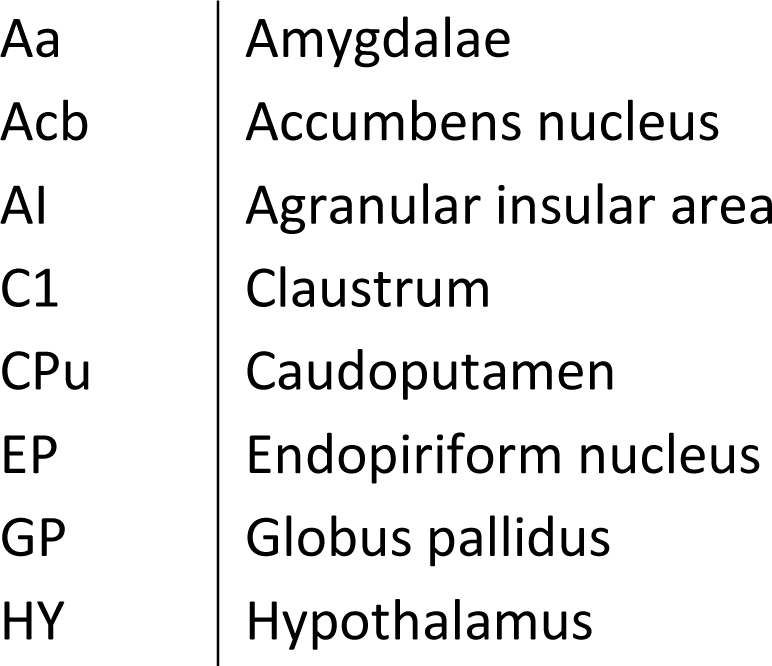
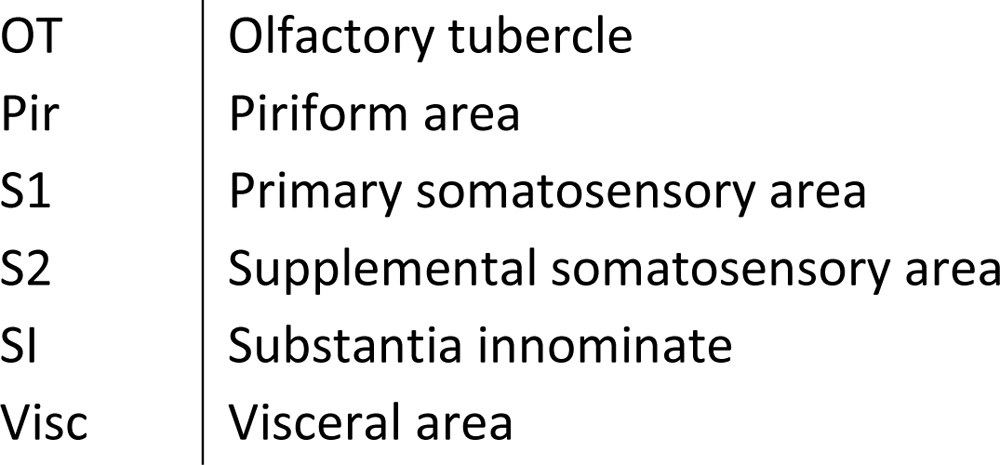
Nomenclature of the brain regions affected by ischemia and corresponding abbreviations

Since iodine stain conferred more contrast to myelin than any other structure, TIA model brains were further processed at higher resolution scans and a semi-automatic segmentation was performed combining manual segmentation of the caudate putamen with the thresholding of white top-hat transformed image (**Fig. 4G**) and a method for quantification (**Fig. 4H,I,J**) of the white matter fibers in 3-D was developed. After 10-minute ischemia, white matter fibers of the ipsilateral hemisphere occupied less 14% of the caudate putamen volume, when compared to the respective contralateral hemisphere (**Fig. 4I**), probably due to fiber decreased wall thickness (**Fig. 4H**). This method allowed a clear discrimination and quantification of the white matter degeneration in the ipsilateral striatum after a 10-minute ischemic insult. Taken together, these results indicate that high resolution ex-vivo micro-CT (contrast-enhanced) constitutes a major advantage to the mouse preclinical stroke studies, improving the time, workload and increasing the outcomes compared with traditional histological methods.

### Comparison of micro-CT imaging of mouse infarct/edema with histology staining

Lesion volumes and edema extent obtained from the 3D reconstructed datasets, obtained with micro-CT (after osmium tetroxide contrast agent enhancement), was correlated with the histological TTC method for lesion quantification (**Fig. 8A**). TTC staining is the gold-standard, or at least the most used methodology for detecting and quantifying infarcts in rodents stroke models ^56^. This method is rather easy to perform and is based on the reduction of the TTC by mitochondrial oxidative enzymes. If the cells in the tissue are alive, reduction of TTC occurs and it turns into a red pigment, while the areas of the tissue with dead cells remain white. However, this method displays important limitations: it can only be performed in fresh thick tissue slices (then fine infarct localization is lost); it only accounts for healthy mitochondria, and does not address other effects such as white matter degeneration, e.g.; can/should only be used in the first 24 h, since after that period occurs the infiltration of inflammatory cells (containing live mitochondria) ^57^. Thus, when comparing sham and tMCAO animals at 3h, and importantly, at 24h after reperfusion, both in edema extent (**Fig. 8C**) and total lesion values (**Fig. 8D**), micro-CT and TTC presented correlation indexes of 0.96 and 0.81 for each parameter, respectively (**Fig. 8C,D**). This means that the lesions volumes obtained from micro-CT reconstructions correlated positively with TTC histopathology. Hence, this novel methodology can be used to accurately determine infarct volumetry in rodent stroke models.

**Fig. 7.**
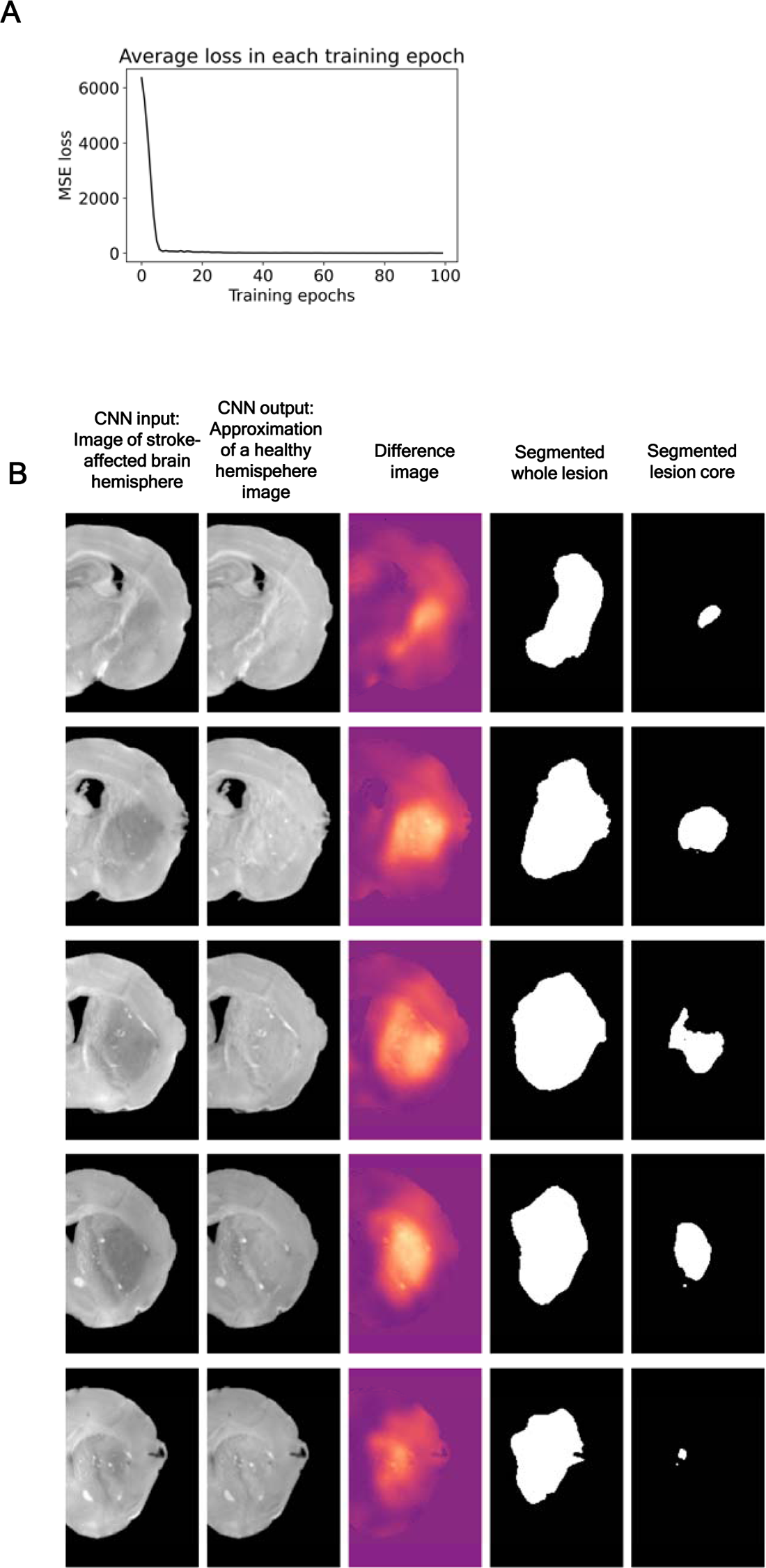
Neural network training and results visualization. In **(A)** the mean training loss (MSE) in each training epoch is plotted. In **(B)** the trained CNN is applied on real lesion brain hemispheres and the segmentation of the whole lesion and lesion core is visualized.

**Fig. 8.**
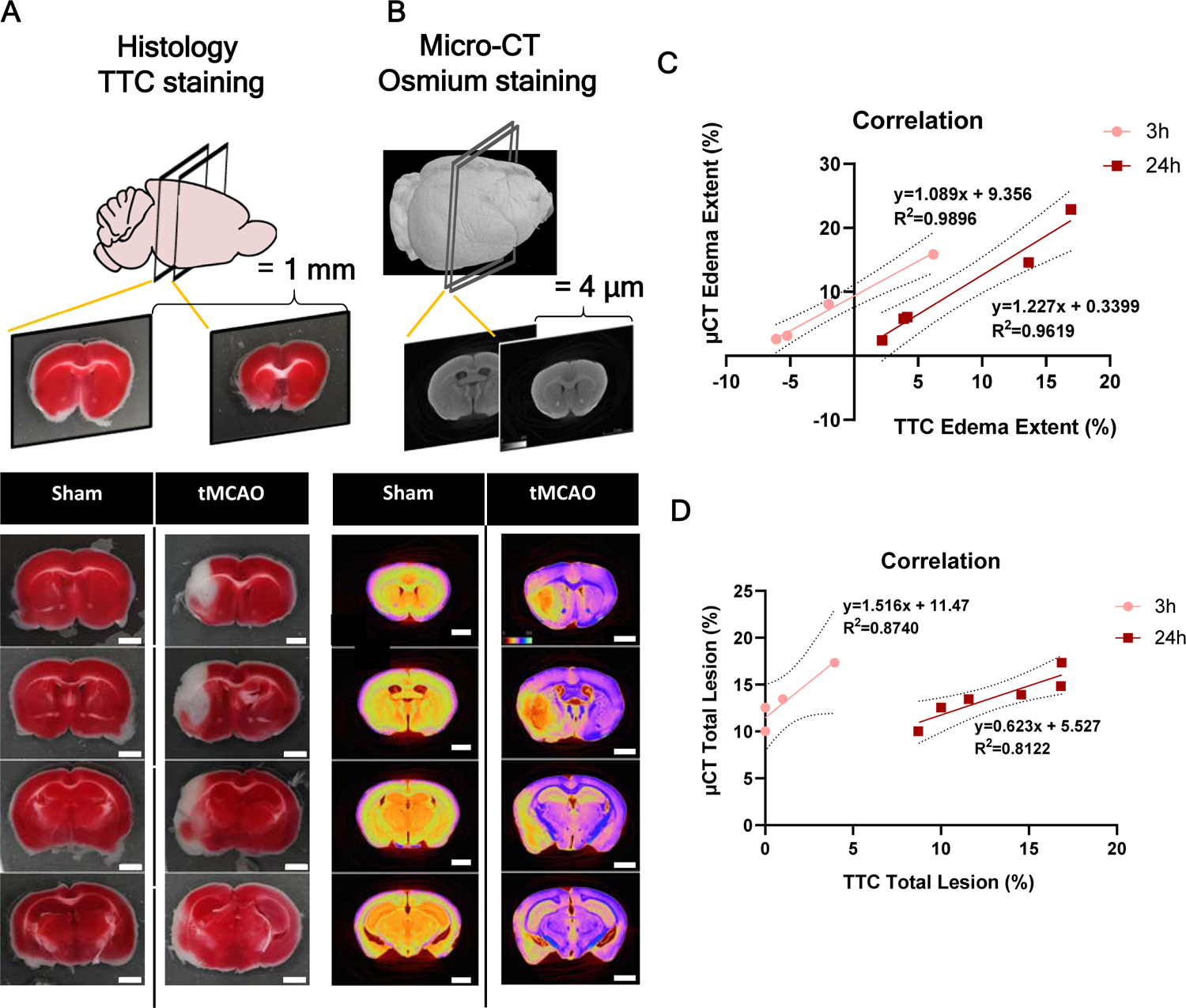
Ex-vivo micro-CT imaging finely correlates with classical histological method TTC, infarct/edema quantification. **(A)** Representative histological cross-sections of TTC stained brains slices of both sham and tMCAO 45-minute mice brains, at 24h-reperfusion. **(B)** Representative 3D micro-CT volume renders cross-sectioning of both sham and tMCAO 45-minute mice brains, at 24h-reperfusion. **(C)** Linear regression of edema extent (%) values measured by histology and micro-CT, at both 3- and 24-hours reperfusion. With the intercept fixed at zero, there is a significant correlation between the two measurement techniques (3h: p=0.0052, R^2^=0.9896; 24h: p=0.0032, R^2^=0.9619; 3-hour reperfusion: micro-CT brains, n=4. TTC brains, n=4. 24-hour reperfusion: micro-CT brains, n=4; TTC staining, n=8). **(D)** Linear regression of total lesion (%) values measured by histology and micro-CT, at both 3- and 24-hours reperfusion. With the intercept fixed at zero, there was significant correlation between the two measurement techniques at 24-hours reperfusion (3h: p=0.0651, R^2^=0.8740; 24h: p=0.014153, R^2^=0.8122; 3-hour reperfusion: micro-CT, n=4; TTC, n=4. 24-hour reperfusion: micro-CT, n=6, TTC, n=6).

### High resolution micro-CT imaging allows data co-registration with mouse brain atlas and identification of specific brain areas affected

Reporting the location of the lesion, its extent, and how consistent these results are among different test subjects is important in preclinical stroke studies. Additionally, lesion registration to a specific sub-anatomical brain structure can further be linked to functional consequences ^19^. Taking advantage of the serial section aligner QuickNII® software/tool that has the mouse (C56BL7) anatomical brain area identified and annotated according to the Allen Mouse Brain Atlas^58 59^, high resolution micro-CT mouse brain sections can be associated to the corresponding brain region. We aligned images of a stroke model, 45min tMCAO mice model (24 h reperfusion), with the serial section aligner QuickNII ® tool (**Fig. 9A-C**), and identified the infarct affected anatomical areas^58 59^ (**Table 2**). The most sensitive anatomical area to ischemia in our model was the caudate putamen, which was altered after ischemic periods as small as 10 minutes. When looking into longer ischemic periods, the infarct consisted usually of the caudate putamen and several functional structures of the cortex involved in motor, sensory and associative circuits, such as the primary somatosensory, the gustatory, the agranular insular and the piriform areas.

**Fig. 9.**
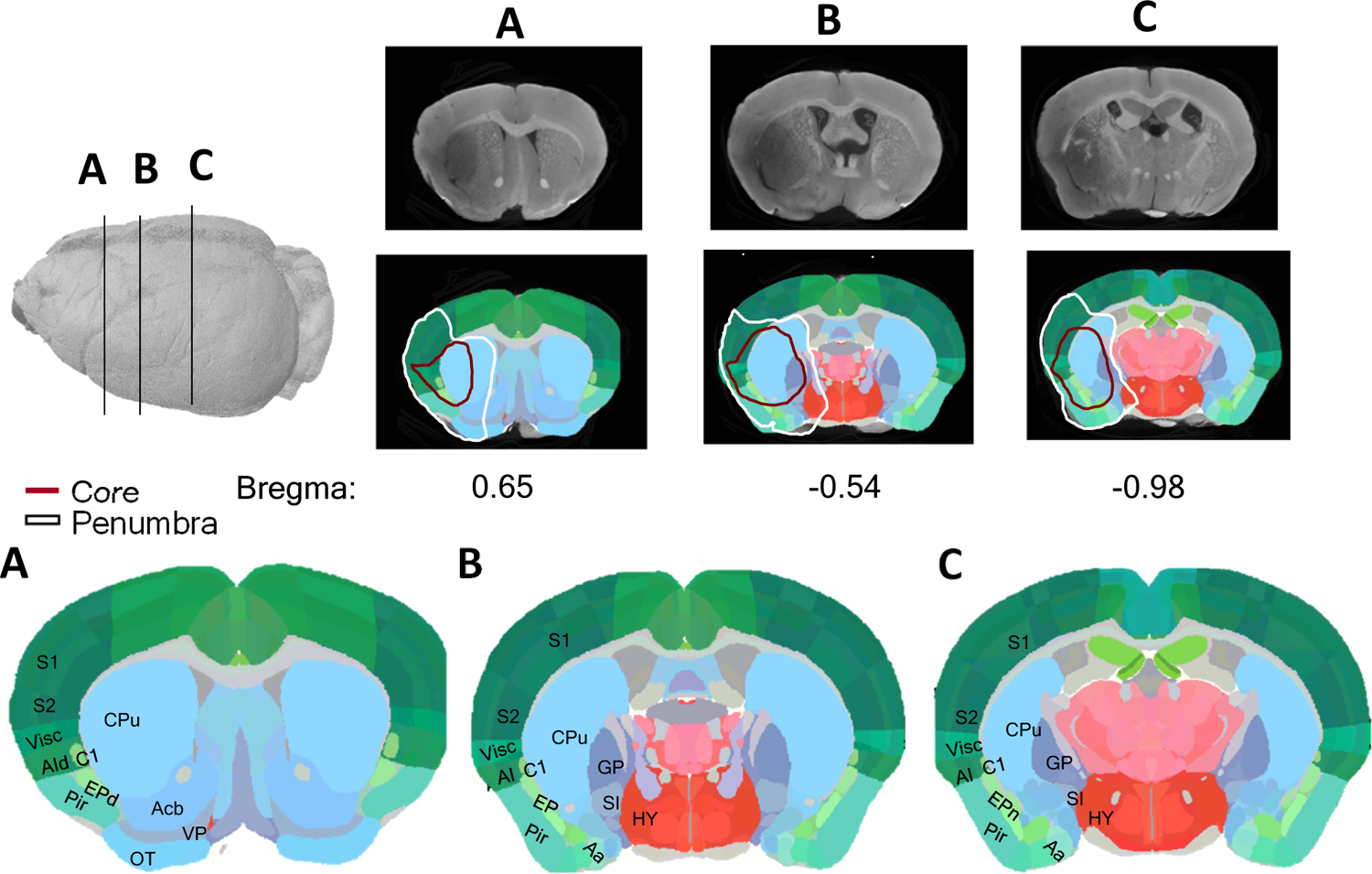
Identification of brain areas by co-registration of micro-CT images with Allen Mouse Brain Atlas, using QuickNII ®. JPEG images of whole 3D volume of a representative tMCAO 45-minute (24h reperfusion) mouse brain were uploaded to QuickNII software, and every 10 sections were aligned with the representative brain atlas. Lesion affected brain structures were assessed and identified according to **Table 3**.

## DISCUSSION

In this work, we report for the first time that micro-CT imaging can be a technique of major advantage for the detection and fine quantification of *ex vivo* brain lesions after ischemia, as well as for assessment of white matter degeneration, in rodent models of stroke. The visualization of the brain’s internal structures by high resolution CT imaging is only possible by using a contrast agent. Considering that, in the literature, several heavy-metal stains have been commonly reported for contrast enhancement of soft tissues, we compared osmium tetroxide, inorganic iodine, iohexol, PMA and PTA for their staining capability, as well as their ability to yield useful anatomical contrast and distinguish established ischemic consequences in the brain of mice subjected to ischemia models. Staining with osmium tetroxide and inorganic iodine yielded the best anatomical contrast in comparison to the other substances tested (iohexol, PTA and PMA (**Supplementary Fig.1**)). Similar staining protocols for PTA and PMA had been used by Descamps, E., et al. ^16^ to stain whole mouse embryos, however in our study these compounds did not diffuse completely into the mice brain, probably due to its bigger size. Dobrivojevic, M., et al. ^60^ used iohexol to identify an ischemic lesion in a tMCAO mice model. However, their data show low contrast on brain anatomic structures and poor ischemic lesion imaging, when compared to what we observed in the present study. On the other hand, osmium tetroxide and the inorganic iodine staining protocols based on Masis, J., et al. ^22^ and Zikmund, T., et al. ^16^, respectively, resulted in contrast enhancement of brain anatomy. Furthermore, they allowed visualization of ischemic degeneration, such as the definition of the lesion area, edema, and white matter degeneration. Since, iodine binds to glycogen molecules, the contrast obtained by this agent reflects the bulk of water in the brain tissue, and because of this myelin and grey matter display higher contrast, while white matter neurons appear more attenuated ^39^. Meanwhile, osmium binds mainly to phospholipids and proteins, reflecting cellular density ^26^. Iodine allowed better discrimination between brain structures due to higher grey/white matter contrast, meanwhile osmium tetroxide still proved to be sufficiently powerful to distinguish and demarcate most of the anatomical brain features. And while both stains allowed the visualization and manual segmentation and quantification of the most remarkable stroke consequences (definition of lesion area and edema), automatic segmentation of white matter fiber tracts and infarct lesion was only possible in iodine-stained brain images. Furthermore, a contrast agent should be chosen also for their manipulation safety and ease of handling besides their effectiveness ^38^. Stains based on inorganic iodine, thus, are easier to handle and much less toxic than osmium tetroxide, besides penetrating faster into the tissue and producing higher contrast in x-ray images, allowing manual/automatic segmentation. Micro-CT imaging matched tissue contrast achieved by longer, more expensive, and step intensive MRI, while delivering higher voxel diameter resolution. Moreover, micro-CT guarantees that the cerebral morphology remains intact and allows the construction of 3D models with higher level of detail and resolution ^61^. However, micro-CT investigation with this resolution in soft tissues can only be performed *ex-vivo* post-mortem, whereas MRI permits longitudinal assessment of lesion growth in the same individual.

Besides, the 5-hour scan is much faster than the ex-vivo micro-MRI scans that have been reported on the literature, where a mouse head scan with isotropic resolution 62.5 µm has been reported to take up to 12h of scanning^62^. Additionally, the 24-hour to 2-week period for sample processing, together with the 3- to 5-hour-long CT measurement can provide a quick and complete visualization of the brain’s external and internal morphology, with optical slices of 3.5-4 µm, considerably thinner than physical slice thickness usually used on histology (0.4 – 1 mm). Since sample preparation for micro-CT only requires solution exchanges, thus it takes roughly the same amount of time to process one sample as it does many samples, making it easier to process a large number of brains simultaneously than in histological methods, where each brain has to be sliced individually ^63^.

Occlusion of the MCA for 45-minutes creates a clearly defined infarct area, usually comprising part of the cerebral cortex and the striatum of the ipsilateral hemisphere, therefore mimicking the location of most common human stroke lesions ^16^. Histological methods for infarct analysis usually comprise brain slicing followed by either enzymatic labelling of live cells (as is the case with TTC, using fresh tissue) or immunolabeling for specific cellular markers (as NeuN or MAP2, using fixed tissue). This might give rise to images with higher-resolution, up to the cellular and subcellular level, but only in the bidimensional context, and with tissues suffering shrinkage and deformation due to processing, besides the possibility for information loss due to slicing ^1^. Infarcted brains, when soaked/stained in either inorganic iodine or osmium tetroxide, allowed the visualization/manual quantification of the total lesion area and edema, as well as the discrimination of the peri-infarcted area, which is still difficult to delineate using histology. Even though iodine-based stains also lead to tissue shrinkage, they still retain the 3D morphological information of the sample ^64^. We validated this quantification method by showing that identified lesion correlated positively with prevalent histopathology observed in TTC processed brains. In addition, the possibility of further using iodine-stained brains for processing using histological procedures, such as immunolabelling, might extend even further the 3D virtual histological results obtained, by precise identification and correlation of different cell types to anatomical structures visualized in the micro-CT^38 46^. Thus, tremendous amount of data can be generated from only one animal, giving rise to gigabytes of information that need very powerful computers for storage, transfer, processing, display, and analysis ^65^.

TIA model is defined by successful occlusion of the MCA, in which mice do not present any neurologic deficit nor lesion (on MRI) at 24-hours reperfusion. This model has not been broadly studied since no morphological alterations have been observed by using TTC staining or MRI and only through cellular/in-situ methods (such as Haematoxylin Eosin stain, and TUNNEL) ischemic alterations can be identified. Nonetheless, the majority of the studies on this model agrees that short term ischemia leads to selective neuronal loss in the striatum of the affected hemisphere and that TIA causes functional deficits in human patients ^66^ ^7^. White matter injury, as characterized by demyelination and axonal breakdown, was also found after MCAO in rats, and this injury played a critical role in these rodents’ poststroke disability ^13^. Segmentation of lesion constitutes a prerequisite for quantitative analysis in stroke research, but this is a tedious and bias/error-prone task when done manually. This process is also time consuming, limiting the sample size that can be analysed at any given time. Thus, the present work addressed the need for reliable, semi-automatic or automatic methods to derive high-quality lesion segmentations and quantifications ^67^. Here, we demonstrate a new method that allows the observation and semi-automatic quantification of white matter degeneration in the striatum after 10-minute tMCAO in mice, allowing for future and more informative studies on this pathology. The utilization of this method was enabled by the high contrast provided by the iodine stain. This allowed us to observe the white matter volume in each brain hemisphere with respect to the caudoputamen volume. Furthermore, we detected a decrease in the white matter volume in the stroke-affected hemisphere. As manual input in the first phase of the segmentation procedure is still required for the segmentation of the striatum in each brain hemisphere, a human-originated error can be introduced into the evaluation. To be objective, the manual segmentation was performed with the Allen Mouse Brain Atlas as a reference. In the following step, we apply thresholding in the both brain hemispheres simultaneously, we can expect the quantitative evaluation of the white matter volume fraction to be consistent between brain hemispheres. The deep learning-based segmentation of the lesion core and penumbra allows for a fully automatic and consistent evaluation of the lesion volume across multiple samples. We have also shown that ground image segmentation masks are not necessary to train and apply the lesion segmentation model and by simulating the lesion on healthy brain hemisphere, we were able to obtain volume measurements that highly correlate with those obtained by manual segmentation. Thus, the uncertainty of manual segmentation is avoided and the data is processed without the time-demanding manual segmentation. The limitation of the method stems from the fact, that a human operator must be able to separate the healthy brain hemisphere from the hemisphere affected by stroke. Furthermore, the lesion must be isolated in one of the brain hemispheres. To lower the computational requirements of the proposed method, the segmentation is performed on images, which were down-sampled 10 times, however, since high-frequency image features are not necessary for the lesion detection, this does not negatively affect the results.

Lesion size and location affects the magnitude of impairment and recovery following stroke^6^. Co-registration of volumetric images obtained with micro-CT, using the QuickNII serial section aligner, based on Allen Mouse Brain Atlas, enables the identification of anatomical brain structures that were affected within the studied mouse models, granting even more functional information. However, this co-registration is still manual and should be considered with caution, due to brain edema on the ipsilateral side. Nonetheless, this overlap with the atlas images allowed the identification of major sub-anatomical brain areas affected in tMCAO model, including the striatum and the somatosensory cortex, among others (**Fig. 9**). As a matter of fact, affected areas can be correlated with the deficits that the mice presented within functional tests and compared to human patient condition^19 59^. Nevertheless, registrations of ischemic lesions in stroke models to brain atlas has only been previously performed on MRI images ^19^, and as mentioned, MRI images are prone to distortion. No such registration was performed using more accurate micro-CT images before our study.

In conclusion, we describe the use of micro-CT as a powerful tool for easy, quick, non-destructive 3D imaging of the whole mouse brain, with high resolution and detail. Iodine- and osmium-stain of mouse brains subjected to different ischemia times allowed the characterization of ischemic lesion evolution overtime. Manual quantification of ischemic lesion, with clear discrimination of the core area (when present), was also possible, with a positive correlation to quantification of the whole lesion by TTC staining. Conversely, only iodine-contrast allowed for automatic quantification of core/penumbra volumes on stroke brains, as well as characterization of striatum white matter degeneration on TIA brains. Therefore, the novel methodology characterized in the present work constitutes a major asset to preclinical stroke studies.

## Material and Methods

### Mice

The number of mice handled for the presented work was approved by the Institutional and National General Veterinary Board Ethical Committees (approval reference number 003424), according to National and European Union rules. Three- to 12-month-old, male/female wild type C57BL6 mice were used. The animals were reproduced, maintained (regular rodents chow and tap water ad libitum) and experimentally manipulated under a 12h light/dark cycle in type II cages in specific pathogen-free conditions in the animal facility (microbiological heath status available). The method of euthanasia used was lethal anesthesia with 170 mg/kg ketamine/ 2 mg/kg medetomidine via ( ketamine + medetomidine, intraperitoneal administration. Charles River Laboratories is the external animal facility used to acquire animals. Physical randomization for group selection was performed using a bioinformatics tool available from Graphpad Prism® QuickCalcs website: https://www.graphpad.com/quickcalcs/randomize1/. The order by which the animals from the different experimental groups were assessed was random during each experimental session. ARRIVE guidelines were taken into consideration in experimental reporting. This study was not pre-registered.

### tMCAO

Mice were anaesthetized with isoflurane (4% for induction and 2-1.25% for maintenance) in a mixture of N_2_O:O_2_ (70:30) using a small-anesthesia system. Rectal temperature was maintained at 37.2°C throughout the surgical procedure making use of a rodent warmer with rectal probe (Stoelting, Wood Dale, IL, USA).

Cerebral blood flow (CBF) was monitored trans cranially using a laser Doppler flowmeter (LDF, PeriFlux System, Perimed, Stockholm, Sweden). Animals were placed under a stereo microscope (Leica S8 APO, Leica Microsystems, Wetzlar, Germany) and fur from the mice’s nape was shaved and disinfected. An incision of 1-2 cm was made on the clean skin, and the margins of the incision were pulled laterally to reveal the cranium. A 0.5 mm diameter microfiber Laser-Doppler probe (Master Probe 418-1connected to microtip: MT B500-0L240) was attached to the skull with cyanoacrylate glue 6 mm lateral and 2 mm posterior to bregma. While under general anesthesia, regional cerebral blood flow was monitored within the MCA territory. The surgical procedure was considered adequate if ≥70% reduction in blood flow occurred immediately after placement of the intraluminal occluding suture and reperfusion occurred after occlusion period; otherwise, mice were excluded.

Transient middle cerebral artery occlusion was induced by the intraluminal suture method^53 68 69 70^. Briefly, after a laser-Doppler probe was attached to the skull, animals were placed in a supine position, under the stereo microscope. The surgical region of the neck was shaved and disinfected. A midline neck incision was made, followed by dissection of the underlying nerves and fascia to expose the right common carotid artery (CCA). The CCA was carefully separated from the vagus nerve, which lies laterally to it, taking extra care not to damage/stimulate or puncture it with the surgical tools. The right external carotid artery (ECA) and internal carotid artery (ICA) were also isolated by blunt dissection of the surrounding tissues (avoiding the rupture of the superior thyroid and occipital arteries). Afterwards, the CCA was temporarily ligated with a silk suture and the ECA was permanently ligated as distal as possible from the bifurcation of the CCA, and loosely ligated proximal to the bifurcation. A reduction in cerebral blood flow was observed. A microvascular clamp was applied to the ICA. A small incision was made in the ECA between the silk sutures, and a new silicone-rubber coated 6-0 nylon monofilament ([21-23] 6023PK10 Doccol Corporation, Massachusetts, USA) was introduced into the ECA and pushed up the ICA until the filament was ≈9 mm from the place of insertion (marked with a silver sharpie©) and a sudden drop was observed in the blood flow, effectively reaching the circle or Willis and blocking the MCA. The coating length of the monofilament lied between 2-3 mm, and the tip diameter depended on the animal’s weight. The loose suture around the ECA was then tightened around the inserted filament to prevent movement during the occlusion period, which lasted for either 45 or 10 minutes of occlusion. After designated occlusion time was over, the filament was removed and the proximal section of the ECA was permanently tied. The temporary tie around the CCA (only closed when necessary to avoid blood leakage through the ECA hole) was removed and reperfusion was established. This was observed in the Laser-Doppler as a rise in the regional CBF. Mice were administered 0.2mL of bupivacaine (2mg/kg in 0.9% NaCl) in the neck open wound. The Laser-Doppler probe was cut close to the skull and both the neck and head incisions were sutured. As a control, sham-operated animals were subjected to the same procedure without occlusion of MCA. Immediately after surgery, a 0.5 mL saline (0.9% NaCl) and buprenorphine (50 µL/ 10 g of animal weight − 0.08 mg/kg) were given subcutaneously to each mouse as fluid replacement and as a form of analgesia, respectively. Buprenorphine was repeatedly administered at time intervals of 8 to 12 hours for the first 3 days post-ischemia; two daily administrations of 0.3 mL of both 5% glucose in 0.9% NaCl and 50/50 % Duphalyte in NaCl were given until mice recovered weight. Right after surgery, mice were placed in a recovery box (Small Animal Recovery Chamber/Warming Cabinet, Harvard Apparatus) maintained at a humidified temperature of 36.8°C until fully recovered from anesthesia (∼2h). To reduce stress, some elements such as paper from the home nest and roll papers were placed in the recovery box. When anesthesia completely wore off, recovery box temperature was decreased to 34.8° for the next 12 hours; afterward, home cages were half placed on a heating pad for the first day post-surgery, allowing mice to choose their environment during recovery, regulating body temperature and controlling post-stroke hypo/hyperthermia. When mice were returned to their home cage, significant care was taken into pairing mice with their pre-surgery cage mates. However, we separated sham-operated animals from tMCAO-operated animals to avoid stress and fighting within the cage, due to differences in the alertness and motile/cognitive deficits between the animals. Mortality rate was 10% (in tMCAO 45minutes only). After 3-, 24- or 72-hours of recovery, animals were euthanized.

### Osmium tetroxide Stain

In the Osmium tetroxide cohort, mice were deeply anesthetized and were perfused transcardially (using 20ml syringes, through high-pressure manual delivery, keeping heart inflated) with 40 mL PBS, followed by 40mL of 2% PFA + 2.5% glutaraldehyde with 4% sucrose in PBS, pH 7.4. The whole brain was removed and fixed in 30 mL of the same fixation solution for at least 1 week, at 4°C, in a 50ml conical centrifuge tube.

After the post-fixation period, brains were washed in water to remove fixatives and PBS, and stained in aqueous 2% osmium tetroxide (O_s_O_4_) solution (15ml for each mouse brain) in 50ml conical centrifuge tube for two weeks, at 4°C in a horizontal shaker. Since osmium tetroxide is toxic if inhaled, tubes were sealed with Parafilm® to minimize evaporation and osmium tetroxide manipulation was always performed in a fume hood. When performing the staining, tubes were protected from direct light with aluminum foil. After staining, brains were washed, wrapped in Parafilm® (to prevent dehydration during micro-CT scanning) and transferred to Parafilm® sealed 5ml tubes (Sarstedt, 62.558.201) for µCT scanning, for short term storage. If long term storage is needed, brain dehydration followed by resin infiltration and embedding should be performed, as described by Masís et al ^71^. In this work, short term storing was the chosen option, mainly because one of the parameters to be analyzed was the brain edema after tMCAO. As such, the dehydration procedure could influence the measurement of this parameter, even though edema is always calculated in comparison with the contralateral hemisphere of the corresponding brain.

Detailed information regarding all the steps involved in *ex-vivo* mouse osmium tetroxide staining is available in Table 4.

**Table 4.** Detailed protocol for whole mice brain preparation for microCT with osmium tetroxide.

| Step | Solution | Time (min) | Temp. (C°) |
| --- | --- | --- | --- |
| **Perfusion** |  |  |  |
| Lethal anesthesia | Ketamine/ medetomidine, IP | 5 | 22 |
| Blood perfusion | PBS | 10 | 22 |
| Brain fixation | 2% PFA, 2.5% GA, 4% sucrose, in PBS | 10 | 22 |
| **Post-fixation** |  |  |  |
| Fixation | 2% PFA, 2.5% GA, 4% sucrose, in PBS | 7-30 days | 4 |
| Washing | Distilled H<sub>2</sub>O | 1, 1, 1, 15 | 22 |
| **Staining** |  |  |  |
| Osmium | 2% OsO<sub>4</sub>, in H<sub>2</sub>O (15 mL/brain) | 15 days | 4 |
| **Micro-CT prep** |  |  |  |
| Washing | Distilled H<sub>2</sub>O | 1, 1, 1, 15, 15, 15 | 22 |
| Preparation | Parafilm® wrapping, and 5 mL_ Tube accommodation | 5 | 22 |
| **Brain Storing** |  |  |  |
| Short-term | Protected from light; Sealed container | 3-12 months | 4 |
| Long-term | Dehydration; Resin embedding | Years | 22 |

### Micro-CT scanning for Osmium stained brains

All osmium tetroxide stained ex-vivo mouse brains were scanned using Bruker micro-CT scanner (SkyScan1276, Bruker, Belgium), from the i3S Scientific Platform Bioimaging. The settings used were: voltage of 90 KV and an output current of 47 µA; 4 µm spatial resolution, 0.2° rotation step through 360 degrees, giving rise to 1801 projections; an Aluminum filter of 1 mm was used, together with a frame averaging of 4 The scanning of each brain took around 5h.

Reconstruction of scanned brain images was performed using the NRecon software (version 1.7.4.2, Bruker, Kontich, Belgium). The settings were normalized for all the brains, however fine tuning was performed for each individual brain to improve quality of deconvolution not achieved with normalized automatic deconvolution. Fine tuning parameters were: Smoothing of 0; Misalignment compensation between 52-60; Ring artifacts reduction between 4-6; and Beam-hardening correction between 20-30%.

### Iodine staining

After the aforementioned recovery times, the animals from the iodine cohort were anesthetized and transcardially perfused (using 20 mL syringes, through manual delivery – keeping heart inflated) with 40 mL PBS, followed by 20 mL of 4% PFA in PBS, pH 7.4. Brains were carefully dissected and post-fixed in 20 mL of the same fixation solution for at least 3 days, at 4°C, in a 50 ml conical centrifuge tube.

Brains were dehydrated in ethanol grade series (30%, 50%, 70%, 80%, 90%) for 2 h followed by a 24-hour staining in 90% methanol plus 1% iodine solution, in agitation, at room temperature. Since iodine solutions are not toxic, no protective measurements were not needed.

After staining, brains were quickly rehydrated with 30% and 70% ethanol, for 1 hour each, at room temperature with agitation. Finally, they were wrapped in Parafilm® (to prevent further dehydration during the micro-CT scan) for micro-CT scanning. For short-term storing, brains were kept in 5 mL tubes (Sarstedt, 62.558.201) at 4°C. If long term storage is needed, brain dehydration, followed by resin infiltration and embedding should be performed, as described by Masís et al ^16^. In this work, short term storage was the chosen option.

Detailed information regarding all the steps involved in *ex-vivo* mouse iodine stain is available in Table 5.

**Table 5.** Detailed protocol for whole mice brain preparation for micro-CT with iodine.

| Step | Solution | Time (min) | Temp. (C°) |
| --- | --- | --- | --- |
| **Perfusion** |  |  |  |
| Lethal anesthesia | Ketamine/ medetomidine, IP | 5 | 22 |
| Blood perfusion | PBS | 10 | 22 |
| Brain fixation | 4% PFA, in PBS | 10 | 22 |
| **Post-fixation** |  |  |  |
| Fixation | 4% PFA, in PBS |  | 4 |
| **Staining** |  |  |  |
| Dehydration | 30%, 50%, 70%, 80%, 90% ethanol | 2 hours, each | 22 |
| Iodine | 1% iodine, in 90% metanol (15 mL/brain) | 24 hours | 22 |
| **micro-CT prep** |  |  |  |
| Preparation | Parafilm® wrapping | 2 | 22 |
| **Brain Storing** |  |  |  |
| Short-term | Protected from light; Sealed container | 3-12 months | 4 |
| Long-term | Dehydration; Resin embedding | Years | 22 |

### Micro-CT scanning for iodine stain

Iodine stained *ex-vivo* mouse brains were scanned using the Bruker micro-CT scanner (SkyScan1276, Bruker, Kontich, Belgium), from the i3S Scientific Platform Bioimaging, where the settings used were: voltage of 85 kV and an output current of 200 µA; 3 µm spatial resolution, 0.2° rotation step through 360°, giving rise to 1801 projections; an Aluminum filter of 1 mm was used, together with a frame averaging of 8. Scanning of each brain took around 3h.

Reconstruction of scanned brain images was performed using the NRecon software (version 1.7.4.2, Bruker, Kontich, Belgium). The settings were normalized for all the brains, however fine-tuning was performed for each individual brain to improve the quality of reconstruction not achieved with normalized automatic deconvolution. Fine-tuning parameters: Smoothing of 0; Misalignment compensation between 52-60; Ring artifacts reduction between 4-6; and Beam-hardening correction between 20-30%.

For higher resolution scans, used for striatum fiber automatic segmentation, stained samples were fixed in 1% agarose gel (Top-Bio s.r.o., Prague, Czech Republic) and placed in 1.5 mL centrifuge tube. GE Phoenix v|tome|x L 240 (Waygate Technologies GmbH, Huerth, Germany) was used for the acquisition. The micro-CT scan was carried out in an air-conditioned cabinet (21 °C) at 60 kV acceleration voltage and 200 μA tube current. Exposure time was 700 ms and 3 X-ray projections were acquired in each angle increment and averaged for reduction of noise. The achieved resolution was 4.5 μm with isotropic voxels. Tomographic reconstruction was performed by using the software GE phoenix datos|x 2.0 (Waygate Technologies GmbH, Huerth, Germany). The data was then imported to VG Studio MAX 3.4 (Volume Graphics GmbH, Heidelberg, Germany).

### Manual segmentation and quantification of total lesion/ core volume and edema using the CTAn software

Fine adjustment of the 3D volume images was performed using the DataViewer© software (version 1.5.6.2, Bruker, Kontich, Belgium), followed by volumetric analysis was performed on CTAn© software (version 1.20.3.0, Bruker, Kontich, Belgium). For edema extent calculation, brain hemispheres were manually annotated in the transaxial orientation every 10 slices. The software rendered the region of interest in every slice though interpolation and calculated object volume. In tMCAO 45-minutes brains, total lesion and core, when present, were manually delineated in the transaxial orientation of every 10 slices. Once again, the software adjusted the region of interest in every slice though interpolation and calculated object volume. Edema extent was calculated by applying the equation: Edema extent (% of the contralateral hemisphere) = ((V _ipsilateral_ (µm^3^)- V _contralateral_ (µm^3^))/ V _contralateral_ (µm^3^))*100; and lesion volume was corrected for edema by applying the equation: Total lesion (% of brain volume) = (V _lesion_ (µm^3^) x (V _Contralateral_ (µm^3^)/ V _Ipsilateral_ (µm^3^)*100)/ V _brain_ (µm^3^); with V being the volume obtained during image quantification and V _brain_ as the sum of V _contralateral_ and V_ipsilateral_, based on ^16^.

### Semi-automatic segmentation and quantification of white matter degeneration in the striatum

The first step required for the analysis of the white matter fibers was their segmentation in the whole CT volume. This segmentation was performed using the software Avizo 2020.2 (Thermo Fisher Scientific, Waltham, MA, USA). The tomographic slices were pre-processed by utilizing the non-local means algorithm for the reduction of noise. Then the caudate putamen was manually segmented in coronal tomographic slices utilizing the coronal Allen Mouse Brain Atlas as reference ^72^. Every 10^th^ slice was manually annotated, and the in-between slices were computed by linear interpolation. These manually segmented volumes were used as a region of interest for subsequent segmentation of the white matter fibers. A semi-automatic approach was used for segmentation of the whiter matter: white top-hat transform. White top-hat transform detects light areas in the image using morphological operators. We utilized a ball-shaped structuring element 19 voxels in diameter. The top-hat-transformed image was then manually thresholded. This segmentation was performed simultaneously for both hemispheres, which allows accurate comparison. For the semi-quantification, a simple volume fraction analysis was performed in the Avizo 2020.2 (Thermo Fisher Scientific, Waltham, MA, USA), where the volume of the white matter was measured in relation with the volume of the manually segmented caudate putamen. The segmented striatum masks were then imported to VG Studio MAX 3.4 (Volume Graphics GmbH, Heidelberg, Germany) and wall thickness analysis was performed. For each voxel, the largest sphere inscribed to the white matter volume was determined, as containing the center position of the evaluated voxel. **See Fig. 6 A** for the overview of the proposed workflow.

### Automatic segmentation of the total lesion and core

An unsupervised deep learning-based approach for the segmentation of the total lesion and core volume was utilized for the automatic evaluation (**See Fig. 6 B)**. For the method to be applied, two conditions must be met: 1) the lesion should be isolated in one brain hemisphere, 2) the user must be able to distinguish the affected hemisphere from the healthy one. First, the brain midline is manually detected, and the brain scans are separated into two subvolumes along this midline. The subvolumes are all cropped to be of unified size of 1920 × 1280 pixels and downsampled 10 times to lower the computational requirements. The hemispheres unaffected by stroke form the training database for the deep learning model. We use a U-net-shaped CNN ^73^ (**See Fig. 6 C** for the schematic representation of the CNN utilized in our work). The size of the CNN input and output is 192 × 128 pixels. The same as in the original U-net implementation, the CNN consists of an encoding and a decoding part. We use 3 × 3 convolutional kernels followed by 3 × 3 strided convolutions (stride 2) in the encoder to perform the repeated down-sampling of the extracted feature maps instead of the max pooling performed in the original implementation. In the decoding part, we apply 3 × 3 convolutional kernels and the up-sampling is performed by 3 × 3 transpose convolution layers with stride 2. The number of convolutional kernels in the first layer of the CNN is 64 and increases up to 1024 in the deepest part of the network. The feature maps from the encoder are concatenated with the decoder feature maps to preserve the spatial information from the input images during the training and inference. ReLU activation function is applied after each convolution. The input to the neural network is a coronal 2-D micro-CT cross-section of the brain hemisphere, where a lesion-like area is randomly simulated by subtracting two randomly generated concentric discs deformed by the elastic transform proposed in ^74^ (parameter alpha is set to 120 and sigma to 5) and blurred with by gaussian blurring with random parameter sigma (range from 5 to 20). The outermost ring simulates the whole lesion and the inner ring simulates the lesion core. The radius of the outer circle is defined randomly in the range from 20 to 80 pixels and its center is randomly set anywhere within the image. The radius of the inner circle is also defined randomly in the range from 0% to 50% of the outer circle radius. The decrease in intensity in the simulated lesion is random from 0% to 50% of the original intensity and is decreased further by up to 50% in the simulated lesion core.

The target of the CNN training is the original CT image of the healthy hemisphere without the simulated lesion area. The CNN thus learns to transform the simulated lesioned brain image into an approximation of images of a healthy brain. The CNN is trained on a total of 830 micro-CT images with randomly simulated lesion for 100 epochs with batch size 32 using the mean-squared-error loss and Adam optimizer ^75^ with initial learning rate of 0.001 (see **Fig. 7 A**). After the network is trained, it can be applied to real lesioned hemisphere image data. The CNN transforms the real 2D micro-CT cross-section of the stroke-affected brain hemisphere into an approximation of healthy brain hemisphere. By subtracting the lesioned image from the output of the CNN, lesion probability map is obtained. This probability map is then automatically thresholded in whole 3D volume by Li thresholding ^76^ to obtain the whole lesion area in 3D (see **Fig. 7 B** for visualization of the segmentation in selected tomographic cross-sections). Threshold for the lesion core is obtained as:

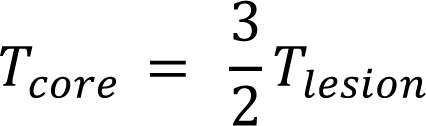

The largest connected component is selected for both the whole lesion area and lesion core thresholded image. The 3D segmented volumes are finally upsampled 10 times by bi-cubic interpolation and form the final segmentation masks and total lesion and core volume can then be computed. Nvidia Quadro P5000 with 16 GB of graphical memory was utilized for training the CNN on a system equipped with 512 GB of RAM and Intel® Xeon® Gold 6248R CPU. The proposed CNN was implemented in the programming language Python (version 3.7.9) using the library Keras ^77^ (version 2.3.1) and Tensorflow backend ^78^ (version 2.1.0). CUDA (version 10.1) and CUDnn (version 7.6.5) were used for GPU acceleration of the training and inference process. NumPy ^79^, scikit-image ^80^ and Pillow libraries were used for manipulating and transforming the image data.

### TTC Stain

In the TTC cohort, infarct volume was evaluated as previously described in ^58^ with minor changes. After the mice were deeply anesthetized, the brain was removed, and the forebrain sliced into 1-mm-thick sections using a mouse brain slicer (Acrylic brain Matrix, Coronal, 40-75 g, Stoelting, USA) on ice. The sections were rinsed once in ice-cold 0.9% sodium chloride (NaCl) for 10 min and subsequently immersed in 25 mL of 0.01% 2,3,5-triphenyltetrazolium chloride (TTC) (Sigma-Aldrich, USA) in 0.9% NaCl at 37°C for 15 min. Slice images were acquired with a camera and analysed *à posteriori*. The unstained area in the section was regarded as an infarct area whereas the stained area was considered non-infarct area.

Infarct volume was determined using a semi-automated quantification method in Fiji-ImageJ ^81^. First, images were calibrated using a millimetric piece of paper as a reference, captured in all photos next to the slices. Next, images were batch processed and analyzed in a custom-made macro. Each image containing several slices of the same condition was pre-processed to isolate each slice, consisting in the manual selection of a colour threshold, followed by a Gaussian filter (sigma 10), thresholded with Huang’s method, and the morphometric operations fill holes and dilation. The output of the preprocessing step is the centroid of each slice. Finally, each slice was segmented alone by duplicating a rectangle, centered in the detected centroids, with size *w x h* (*w* = original image width, *h* = original image height/number of slices).

The infarct lesion measurement of each slice was performed by a blind experimenter with the following steps: i) manual delineation of the middle line to divide the slice into two hemispheres; ii) manual annotation of the stroke area (when present); iii) quality control of the resulting regions of interest. The macro output obtained was a results table with slice labels, the contra- and ipsilateral hemispheres area, and the infarct lesion area. Volumes were obtained by summing the area of the infarct brain segment from all brain slices (considering that each slice’s thickness is 1 mm). Edema extent and corrected lesion volume were calculated as described above and based on ^82^.

### Image registration to Allen Mouse Brain Atlas

The whole image series from a tMCAO mice brain (stained with osmium) were registered to match the cutting plan and proportions of the atlas sections using the open-source serial aligner software QuickNII ^83^ (https://www.nitrc.org/projects/quicknii/). This software allows the assignment of spatial location (features such as angle, and size can be adjusted) so that atlas can match with section images. After, the whole image series segmented with the atlas was saved, and structures affected by ischemic lesion in this mouse were identified.

### Brain slicing and immunolabeling

Brains were post-fixed in the same fixation solution, overnight, at 4°C and then left in 30% sucrose in PBS at 4°C, until they sunk in the solution. Coronal sections (40 µm) were cut on a freezing microtome (Leica Cryostat CM 3050) and used for free-floating immunohistochemistry. Slices were incubated with 10% horse serum, 0.2% Triton X-100 in PBS, for permeabilization and blocking, for 1 hour, in agitation, at room temperature. Primary antibodies were always incubated for 48-72 hours at 4°C, in PBS. Secondary antibodies were incubated overnight at 4°C. The slides were mounted on gelatin-covered glass slides with a fluorescent mounting medium (DAKO) and imaging was performed on a laser scanning Confocal Microscope Leica SP8 (Leica mycrosystems, Wetzlar, Germany), using the ×10 air objective. In each set of experiments, the same batch of antibodies (primary and secondary) were used, and images were taken using the same settings. Primary antibodies used were anti-NeuN (1:500, Mouse, chemicon international, ab377), anti-MAP2 (1:500, Rabbit, abcam, ab246640), and anti-BrdU (TUNEL Assay Kit, Abcam, ab66110); as secondary antibodies Alexa Fluor 488 and 568 (1:500) were used. The fluorescent dye Hoechst 33342 (0.5 μg/mL–15 min room temperature) was used to stain nuclei.

Brains stained with iodine were rehydrated in two, 2-hours, PBS steps and then left in 25% sucrose in PBS at 4°C overnight. Coronal sections were cut and immunolabeled as described above.

## Statistical Analysis

Statistical analysis was performed using GraphPad Prism Version 8.0/9.0 (GraphPad Software, Inc., San Diego, CA, USA). Data were collected from experiments and presented as mean ± SEM with individual points, “n” information (the experimental unit) is described in figure legends. The experimental unit was each single animal. Linear regression analysis was performed to correlate edema extent and total lesion values obtained either through microCT or TTC. For all statistical analysis: ****p<0.0001, ***p<0.001, ** p<0.01, * p<0.05, ns (not significant). Identification and removal of outliers was performed using automatic GraphPad prism software 8.0, using the ROUT (robust nonlinear regression) method, with a Q=1% (recommended).

## Acknowledgements

The authors acknowledge: Maria João Saraiva from i3S for the guidance and laboratory support; Michaela Kavkova from CEITEC for iodine stain protocol details; Andrea Lobo from i3S for reviewing and revising the manuscript for grammar and syntax; Luis Maia from i3S for sharing some reagents and Biostroke project (POCI-01-0145-FEDER-031674) for sharing some mouse stroke micro-CT images, performed by João Gomes (last author of this paper and coPI of the project). The authors also acknowledge the support of the i3S Scientific Platforms: Advanced Light Microscopy (ALM); Bioimaging; and Histology and Electron Microscopy (HEMS), members of the national infrastructure PPBI - Portuguese Platform of Bioimaging (PPBI-POCI-01-0145-FEDER-022122) and the i3S Animal facility.

## Author’s Contribution statement

Raquel Pinto, Jan Matula and João Gomes made a substantial contribution to the concept and design, acquisition and analysis/interpretation of data. Maria G-Lázaro and Tomas Zikmund contributed to acquisition and analysis/interpretation of data. Mafalda Sousa and Josef Kaiser contributed to the analysis/interpretation of data. All authors drafted the article, revised and approved the version to be published.

## Data availability

The data supporting the findings of this study is available within the article, supplementary data or can be available, upon request (due to high Gigabytes of data), from the corresponding author João Gomes.

## Abbreviations

tMCAO: transient Middle Cerebral Artery Occlusion

TIA: transient ischemic attack

MRI: magnetic resonance imaging

micro-CT: microcomputed tomography

PMA: phosphomolybdic acid

PTA: phosphotungstic acid

MCA: middle cerebral artery

CCA: common carotid artery

ECA: external carotid artery

ICA: internal carotid artery

TTC: 2,3,5-Triphenyltetrazolium chloride

## References

1. Balkaya M and Cho S. Optimizing functional outcome endpoints for stroke recovery studies. J Cereb Blood Flow Metab 2019; 39: 2323–2342. 2019/09/17. DOI: 10.1177/0271678X19875212.

2. Murata Y, Rosell A, Scannevin RH, et al. Extension of the thrombolytic time window with minocycline in experimental stroke. Stroke 2008; 39: 3372–3377. 2008/10/18. DOI: 10.1161/STROKEAHA.108.514026.

3. Dirnagl U and Endres M. Found in translation: preclinical stroke research predicts human pathophysiology, clinical phenotypes, and therapeutic outcomes. Stroke 2014; 45: 1510–1518. 2014/03/22. DOI: 10.1161/strokeaha.113.004075.

4. Snyder JM, Hagan CE, Bolon B, et al. 20 - Nervous System. In: Treuting PM, Dintzis SM and Montine KS (eds) Comparative Anatomy and Histology (Second Edition). San Diego: Academic Press, 2018, pp.403-444.

5. Sommer CJ. Ischemic stroke: experimental models and reality. Acta Neuropathol 2017; 133: 245–261. 2017/01/09. DOI: 10.1007/s00401-017-1667-0.

6. Durukan Tolvanen A, Tatlisumak E, Pedrono E, et al. TIA model is attainable in Wistar rats by intraluminal occlusion of the MCA for 10min or shorter. Brain Res 2017; 1663: 166–173. 2017/03/16. DOI: 10.1016/j.brainres.2017.03.010.

7. Pedrono E, Durukan A, Strbian D, et al. An optimized mouse model for transient ischemic attack. J Neuropathol Exp Neurol 2010; 69: 188–195. 2010/01/20. DOI: 10.1097/NEN.0b013e3181cd331c.

8. del Zoppo GJ, Sharp FR, Heiss WD, et al. Heterogeneity in the penumbra. J Cereb Blood Flow Metab 2011; 31: 1836–1851. 2011/07/07. DOI: 10.1038/jcbfm.2011.93.

9. Sharp FR, Lu A, Tang Y, et al. Multiple molecular penumbras after focal cerebral ischemia. J Cereb Blood Flow Metab 2000; 20: 1011–1032. 2000/07/25. DOI: 10.1097/00004647-200007000-00001.

10. Astrup J, Siesjo BK and Symon L. Thresholds in cerebral ischemia - the ischemic penumbra. Stroke 1981; 12: 723–725. 1981/11/01. DOI: 10.1161/01.str.12.6.723.

11. Etherton MR, Wu O, Giese AK, et al. White Matter Integrity and Early Outcomes After Acute Ischemic Stroke. Transl Stroke Res 2019; 10: 630–638. 2019/01/30. DOI: 10.1007/s12975-019-0689-4.

12. Jiang X, Pu H, Hu X, et al. A Post-stroke Therapeutic Regimen with Omega-3 Polyunsaturated Fatty Acids that Promotes White Matter Integrity and Beneficial Microglial Responses after Cerebral Ischemia. Transl Stroke Res 2016; 7: 548–561. 2016/10/08. DOI: 10.1007/s12975-016-0502-6.

13. Li M, Zhao Y, Zhan Y, et al. Enhanced white matter reorganization and activated brain glucose metabolism by enriched environment following ischemic stroke: Micro PET/CT and MRI study. Neuropharmacology 2020; 176: 108202. 2020/07/03. DOI: 10.1016/j.neuropharm.2020.108202.

14. Bazan NG, Halabi A, Ertel M, et al. Neuroinflammation. Basic Neurochemistry. 2012, pp.610–620.

15. Sommer CJ. Ischemic stroke: experimental models and reality. Acta Neuropathologica 2017; 133: 245–261. DOI: 10.1007/s00401-017-1667-0.

16. Masis J, Mankus D, Wolff SBE, et al. A Micro-CT-based Method for Characterizing Lesions and Locating Electrodes in Small Animal Brains. J Vis Exp 2018 2018/11/27. DOI: 10.3791/58585.

17. Bernard R, Balkaya M and Rex A. Behavioral Testing in Rodent Models of Stroke, Part I. Rodent Models of Stroke. 2016, pp.199–223.

18. Dorr AE, Lerch JP, Spring S, et al. High resolution three-dimensional brain atlas using an average magnetic resonance image of 40 adult C57Bl/6J mice. Neuroimage 2008; 42: 60–69. 2008/05/27. DOI: 10.1016/j.neuroimage.2008.03.037.

19. Jeffers MS, Touvykine B, Ripley A, et al. Poststroke Impairment and Recovery Are Predicted by Task-Specific Regionalization of Injury. J Neurosci 2020; 40: 6082–6097. 2020/07/02. DOI: 10.1523/JNEUROSCI.0057-20.2020.

20. Boyd LA, Hayward KS, Ward NS, et al. Biomarkers of stroke recovery: Consensus-based core recommendations from the Stroke Recovery and Rehabilitation Roundtable. Int J Stroke 2017; 12: 480–493. 2017/07/13. DOI: 10.1177/1747493017714176.

21. Balkaya M, Kröber JM, Rex A, et al. Assessing post-stroke behavior in mouse models of focal ischemia. Journal of Cerebral Blood Flow and Metabolism 2013; 33: 330–338. DOI: 10.1038/jcbfm.2012.185.

22. Dobrivojevic M, Bohacek I, Erjavec I, et al. Computed microtomography visualization and quantification of mouse ischemic brain lesion by nonionic radio contrast agents. Croat Med J 2013; 54: 3–11. 2013/02/28. DOI: 10.3325/cmj.2013.54.3.

23. Kastner DB, Kharazia V, Nevers R, et al. Scalable method for micro-CT analysis enables large scale quantitative characterization of brain lesions and implants. Sci Rep 2020; 10: 20851. 2020/12/02. DOI: 10.1038/s41598-020-77796-3.

24. Powers WJ, Rabinstein AA, Ackerson T, et al. Guidelines for the Early Management of Patients With Acute Ischemic Stroke: 2019 Update to the 2018 Guidelines for the Early Management of Acute Ischemic Stroke: A Guideline for Healthcare Professionals From the American Heart Association/American Stroke Association. Stroke 2019; 50: e344–e418. DOI: 10.1161/STR.0000000000000211.

25. Deb P, Sharma S and Hassan KM. Pathophysiologic mechanisms of acute ischemic stroke: An overview with emphasis on therapeutic significance beyond thrombolysis. Pathophysiology 2010; 17: 197–218. 2010/01/16. DOI: 10.1016/j.pathophys.2009.12.001.

26. Saito S and Murase K. Ex vivo imaging of mouse brain using micro-CT with non-ionic iodinated contrast agent: a comparison with myelin staining. Br J Radiol 2012; 85: e973–978. 2012/06/08. DOI: 10.1259/bjr/13040401.

27. Mizutani R, Saiga R, Ohtsuka M, et al. Three-dimensional X-ray visualization of axonal tracts in mouse brain hemisphere. Sci Rep 2016; 6: 35061. 2016/10/12. DOI: 10.1038/srep35061.

28. Parlanti P, Cappello V, Brun F, et al. Size and specimen-dependent strategy for x-ray micro-ct and tem correlative analysis of nervous system samples. Scientific Reports 2017; 7: 2858. DOI: 10.1038/s41598-017-02998-1.

29. Ding Y, Vanselow DJ, Yakovlev MA, et al. Computational 3D histological phenotyping of whole zebrafish by X-ray histotomography. eLife 2019; 8: e44898. DOI: 10.7554/eLife.44898.

30. Clark DP and Badea CT. Micro-CT of rodents: state-of-the-art and future perspectives. Phys Med 2014; 30: 619–634. 2014/06/30. DOI: 10.1016/j.ejmp.2014.05.011.

31. Ghanavati S, Yu LX, Lerch JP, et al. A perfusion procedure for imaging of the mouse cerebral vasculature by X-ray micro-CT. J Neurosci Methods 2014; 221: 70–77. 2013/09/24. DOI: 10.1016/j.jneumeth.2013.09.002.

32. Dullin C, Ufartes R, Larsson E, et al. muCT of ex-vivo stained mouse hearts and embryos enables a precise match between 3D virtual histology, classical histology and immunochemistry. PLoS One 2017; 12: e0170597. 2017/02/09. DOI: 10.1371/journal.pone.0170597.

33. Hong SH, Herman AM, Stephenson JM, et al. Development of barium-based low viscosity contrast agents for micro CT vascular casting: Application to 3D visualization of the adult mouse cerebrovasculature. J Neurosci Res 2020; 98: 312–324. 2019/10/21. DOI: 10.1002/jnr.24539.

34. Hlushchuk R, Zubler C, Barre S, et al. Cutting-edge microangio-CT: new dimensions in vascular imaging and kidney morphometry. Am J Physiol Renal Physiol 2018; 314: F493–F499. 2017/11/24. DOI: 10.1152/ajprenal.00099.2017.

35. Schaad L, Hlushchuk R, Barre S, et al. Correlative Imaging of the Murine Hind Limb Vasculature and Muscle Tissue by MicroCT and Light Microscopy. Sci Rep 2017; 7: 41842. 2017/02/09. DOI: 10.1038/srep41842.

36. Quintana DD, Lewis SE, Anantula Y, et al. The cerebral angiome: High resolution MicroCT imaging of the whole brain cerebrovasculature in female and male mice. Neuroimage 2019; 202: 116109. 2019/08/26. DOI: 10.1016/j.neuroimage.2019.116109.

37. Udagawa S, Miyara K, Takekata H, et al. Investigation on the validity of 3D micro-CT imaging in the fish brain. J Neurosci Methods 2019; 328: 108416. 2019/09/01. DOI: 10.1016/j.jneumeth.2019.108416.

38. de Crespigny A, Bou-Reslan H, Nishimura MC, et al. 3D micro-CT imaging of the postmortem brain. J Neurosci Methods 2008; 171: 207–213. 2008/05/09. DOI: 10.1016/j.jneumeth.2008.03.006.

39. Zikmund T, Novotná M, Kavková M, et al. High-contrast differentiation resolution 3D imaging of rodent brain by X-ray computed microtomography. Journal of Instrumentation 2018; 13: C02039–C02039. DOI: 10.1088/1748-0221/13/02/c02039.

40. Prajapati SI, Kilcoyne A, Samano AK, et al. Erratum to: MicroCT-Based Virtual Histology Evaluation of Preclinical Medulloblastoma. Mol Imaging Biol 2017; 19: 483. 2017/04/09. DOI: 10.1007/s11307-017-1079-5.

41. Girard R, Zeineddine HA, Orsbon C, et al. Micro-computed tomography in murine models of cerebral cavernous malformations as a paradigm for brain disease. J Neurosci Methods 2016; 271: 14–24. 2016/06/28. DOI: 10.1016/j.jneumeth.2016.06.021.

42. Kavkova M, Zikmund T, Kala A, et al. Contrast enhanced X-ray computed tomography imaging of amyloid plaques in Alzheimer disease rat model on lab based micro CT system. Sci Rep 2021; 11: 5999. 2021/03/18. DOI: 10.1038/s41598-021-84579-x.

43. Depannemaecker D, Santos LEC, de Almeida AG, et al. Gold Nanoparticles for X-ray Microtomography of Neurons. ACS Chem Neurosci 2019; 10: 3404–3408. 2019/07/06. DOI: 10.1021/acschemneuro.9b00290.

44. Chin A-L, Yang S-M, Chen H-H, et al. A synchrotron X-ray imaging strategy to map large animal brains. Chinese Journal of Physics 2020; 65: 24–32. DOI: https://doi.org/10.1016/j.cjph.2020.01.010.

45. Luo Y, Yin X, Shi S, et al. Non-destructive 3D Microtomography of Cerebral Angioarchitecture Changes Following Ischemic Stroke in Rats Using Synchrotron Radiation. 2019; 13. Original Research. DOI: 10.3389/fnana.2019.00005.

46. Hayasaka N, Nagai N, Kawao N, et al. In vivo diagnostic imaging using micro-CT: sequential and comparative evaluation of rodent models for hepatic/brain ischemia and stroke. PLoS One 2012; 7: e32342. 2012/03/03. DOI: 10.1371/journal.pone.0032342.

47. Park JY, Lee SK, Kim JY, et al. A new micro-computed tomography-based high-resolution blood-brain barrier imaging technique to study ischemic stroke. Stroke 2014; 45: 2480–2484. 2014/07/12. DOI: 10.1161/STROKEAHA.114.006297.

48. Topperwien M, Doeppner TR, Zechmeister B, et al. Multiscale x-ray phase-contrast tomography in a mouse model of transient focal cerebral ischemia. Biomed Opt Express 2019; 10: 92–103. 2019/02/19. DOI: 10.1364/BOE.10.000092.

49. Ronneberger O, Fischer P and Brox T. U-Net: Convolutional Networks for Biomedical Image Segmentation. In: Medical Image Computing and Computer-Assisted Intervention – MICCAI 2015 (eds Navab N, Hornegger J, Wells WM, et al.), Cham, 2015// 2015, pp.234–241. Springer International Publishing.

50. Toulkeridou E, Gutierrez CE, Baum D, et al. Automated segmentation of insect anatomy from micro-CT images using deep learning. bioRxiv 2021.

51. Léger J, Leyssens L, De Vleeschouwer C, et al. Deep Learning-Based Segmentation of Mineralized Cartilage and Bone in High-Resolution Micro-CT Images. Lecture Notes in Computational Vision and Biomechanics. Springer International Publishing, 2020, p. 158–170.

52. Rytky SJO, Huang L, Tanska P, et al. Automated analysis of rabbit knee calcified cartilage morphology using micro-computed tomography and deep learning. 2021; 239: 251–263. DOI: https://doi.org/10.1111/joa.13435.

53. Koch S, Mueller S, Foddis M, et al. Atlas registration for edema-corrected MRI lesion volume in mouse stroke models. J Cereb Blood Flow Metab 2019; 39: 313–323. 2017/08/23. DOI: 10.1177/0271678X17726635.

54. Metscher BD. MicroCT for comparative morphology: simple staining methods allow high-contrast 3D imaging of diverse non-mineralized animal tissues. BMC Physiology 2009; 9: 11. DOI: 10.1186/1472-6793-9-11.

55. Parlanti P, Cappello V, Brun F, et al. Size and specimen-dependent strategy for x-ray micro-ct and tem correlative analysis of nervous system samples. Sci Rep 2017; 7: 2858. 2017/06/08. DOI: 10.1038/s41598-017-02998-1.

56. Bederson JB, Pitts LH, Germano SM, et al. Evaluation of 2,3,5-triphenyltetrazolium chloride as a stain for detection and quantification of experimental cerebral infarction in rats. Stroke 1986; 17: 1304-1308. 1986/11/01. DOI: 10.1161/01.str.17.6.1304.

57. Liszczak TM, Hedley-Whyte ET, Adams JF, et al. Limitations of tetrazolium salts in delineating infarcted brain. Acta Neuropathol 1984; 65: 150–157. 1984/01/01. DOI: 10.1007/BF00690469.

58. Allen. ALLEN Mouse Brain Atlas. Gene Expression 2007: 1–9.

59. Mrzilkova J, Patzelt M, Gallina P, et al. Imaging of Mouse Brain Fixated in Ethanol in Micro-CT. Biomed Res Int 2019; 2019: 2054262. 2019/08/09. DOI: 10.1155/2019/2054262.

60. Descamps E, Sochacka A, De Kegel B, et al. Soft tissue discrimination with contrast agents using micro-CT scanning. Belg J Zool 2014; 144: 20–40.

61. Rodrigues PV, Tostes K, Bosque BP, et al. Illuminating the Brain With X-Rays: Contributions and Future Perspectives of High-Resolution Microtomography to Neuroscience. Front Neurosci 2021; 15: 627994. 2021/04/06. DOI: 10.3389/fnins.2021.627994.

62. Llambrich S, Wouters J, Himmelreich U, et al. ViceCT and whiceCT for simultaneous high-resolution visualization of craniofacial, brain and ventricular anatomy from micro-computed tomography. Sci Rep 2020; 10: 18772. 2020/11/01. DOI: 10.1038/s41598-020-75720-3.

63. Chen K-C, Arad A, Song Z-M, et al. High-definition neural visualization of rodent brain using micro-CT scanning and non-local-means processing. BMC Med Imaging 2018; 18: 38–38. DOI: 10.1186/s12880-018-0280-6.

64. Anderson R and Maga AM. A Novel Procedure for Rapid Imaging of Adult Mouse Brains with MicroCT Using Iodine-Based Contrast. PLoS One 2015; 10: e0142974. 2015/11/17. DOI: 10.1371/journal.pone.0142974.

65. Dullin C, Ufartes R, Larsson E, et al. μCT of ex-vivo stained mouse hearts and embryos enables a precise match between 3D virtual histology, classical histology and immunochemistry. PLOS ONE 2017; 12: e0170597. DOI: 10.1371/journal.pone.0170597.

66. Schaad L, Hlushchuk R, Barré S, et al. Correlative Imaging of the Murine Hind Limb Vasculature and Muscle Tissue by MicroCT and Light Microscopy. Scientific Reports 2017; 7: 41842. DOI: 10.1038/srep41842.

67. Schoppe O, Pan C, Coronel J, et al. Deep learning-enabled multi-organ segmentation in whole-body mouse scans. Nature Communications 2020; 11: 5626. DOI: 10.1038/s41467-020-19449-7.

68. Koizumi J-i, Yoshida Y, Nakazawa T, et al. Experimental studies of ischemic brain edema. Nosotchu 1986; 8: 1–8. DOI: 10.3995/jstroke.8.1.

69. Longa EZ, Weinstein PR, Carlson S, et al. Reversible middle cerebral artery occlusion without craniectomy in rats. Stroke 1989; 20: 84–91. DOI: 10.1161/01.STR.20.1.84.

70. Chan PH, Kamii H, Yang G, et al. Brain infarction is not reduced in SOD-1 transgenic mice after a permanent focal cerebral ischemia. NeuroReport 1993; 5: 293–296. DOI: 10.1097/00001756-199312000-00028.

71. Yang G, Chan PH, Chen J, et al. Human copper-zinc superoxide dismutase transgenic mice are highly resistant to reperfusion injury after focal cerebral ischemia. Stroke 1994; 25: 165–170. DOI: 10.1161/01.STR.25.1.165.

72. Kuts R, Frank D, Gruenbaum BF, et al. A Novel Method for Assessing Cerebral Edema, Infarcted Zone and Blood-Brain Barrier Breakdown in a Single Post-stroke Rodent Brain. Frontiers in neuroscience 2019; 13: 1105–1105. DOI: 10.3389/fnins.2019.01105.

73. Ronneberger O, Fischer P and Brox T. U-Net: Convolutional Networks for Biomedical Image Segmentation. In: Cham %@ 978-3-319-24574-4, 2015, pp.234–241. Springer International Publishing.

74. Patrice YS, Dave S and John CP. Best Practices for Convolutional Neural Networks Applied to Visual Document Analysis %@ 0769519601. Proceedings of the Seventh International Conference on Document Analysis and Recognition - Volume 2. IEEE Computer Society, 2003, p. 958.

75. Kingma DP and Ba J. Adam: A method for stochastic optimization. arXiv preprint arXiv:14126980 2014.

76. Li CH and Lee CK. Minimum cross entropy thresholding. Pattern Recognition 1993; 26: 617–625. DOI: https://doi.org/10.1016/0031-3203(93)90115-D.

77. Chollet F and others. Keras: The Python Deep Learning library. 2018, p.ascl:1806.1022.

78. Abadi M, Barham P, Chen J, et al. Tensorflow: A system for large-scale machine learning. In: 12th {USENIX} symposium on operating systems design and implementation ({OSDI} 16) 2016, pp.265–283.

79. Harris CR, Millman KJ, van der Walt SJ, et al. Array programming with NumPy. Nature 2020; 585: 357–362. DOI: 10.1038/s41586-020-2649-2.

80. van der Walt S, Schonberger JL, Nunez-Iglesias J, et al. scikit-image: image processing in Python. PeerJ 2014; 2: e453. 2014/07/16. DOI: 10.7717/peerj.453.

81. Gomes JR, Lobo AC, Melo CV, et al. Cleavage of the vesicular GABA transporter under excitotoxic conditions is followed by accumulation of the truncated transporter in nonsynaptic sites. Journal of Neuroscience 2011; 31: 4622–4635. DOI: 10.1523/JNEUROSCI.3541-10.2011.

82. Schindelin J, Arganda-Carreras I, Frise E, et al. Fiji: an open-source platform for biological-image analysis. Nature Methods 2012; 9: 676–682. DOI: 10.1038/nmeth.2019.

83. Puchades MA, Csucs G, Ledergerber D, et al. Spatial registration of serial microscopic brain images to three-dimensional reference atlases with the QuickNII tool. PLoS One 2019; 14: e0216796. 2019/05/30. DOI: 10.1371/journal.pone.0216796.

