## Supplemental material for "High resolution micro-CT imaging in mice stroke models: from 3D detailed infarct characterization to automatic area segmentation"

**Supplemental figure legends:**

**Suppl. Fig. 1- Iohexol, phosphotungstic acid (PTA) and phosphomolybdic acid (PMA) staining of whole mice brain.** Representative images of iohexol **(A)** and PTA **(B)** stained whole brains of mice subjected to tMCAO for 45 min following 24 h reperfusion, and PMA **(C)** stained mice brain not subjected to surgery. Iohexol does not confer contrast enhancement to brain structures, whereas PTA and PMA do not penetrate the whole sample/tissue.

**Suppl. Fig. 2- Iodine staining of mouse brain allows further immunostaining analysis. (A)** Representative 3D visualization of iodine-staining mouse brain whole volume (darker red) in relation to its volume prior to fixation, unstained volume. Calculated volumes in CTAn demonstrated that the brain shrunk to a third of its size. **(B)** Images representing a iodine-stained tMCAO brain sliced after micro-CT scan, and labelled with the specific neuronal marker, NeuN and Hoechst 3342 from nuclear stain. Fluorescent specific labelling was observed, and was distinct between infarct lesion and healthy tissue.

**Suppl. Fig. 3- High resolution micro-CT imaging in mouse stroke model (tMCAO 45') using iodine staining. (A)** Virtual slices (~4µm voxels) at arbitrary orientations from micro-CT scan of a stroke mouse brain stained by inorganic iodine showing total ischemic lesion as well as core (orange/darker) and penumbra ((yellow/grey around core) differentiation. Below panel shows lesion detail in the transaxial and sagittal pane. Scale bar: 2 mm.

**Suppl. Fig. 4- High resolution micro-CT imaging in TIA mouse model using Osmium Tetroxide staining. (A)** Virtual slices (~4µm voxels) at arbitrary orientations from micro-CT scan of a TIA brain stained with osmium tetroxide showing degeneration of striatum white fiber matter, indicated by the white dotted surrounded region. Below panel shows lesion detail in the transaxial and sagittal pane. Scale bar: 2mm.

**Suppl. Fig. 5- Brain ischemic lesions (stroke and TIA models) evolution using high resolution micro-CT.** Representative images of mice sham, TIA and stroke mouse models using micro-CT imaging. Sham brains hemispheres mirror each other and demonstrate no ischemic alteration, whereas brains of TIA subjected mice display striatum white matter degeneration. In stroke model (tMCAO 45') mice brain is possible to observe cortical and striatal lesion, as well as to distinguish between core and penumbra at 24 h following reperfusion. Stroke mouse brain model show an increase in lesion size over the tested 72-hour reperfusion period.

**Suppl. Videos:**

**Suppl. Vid. 1- 3D whole brain of mouse stroke model (tMCAO 45') after osmium tetroxide staining and micro-CT imaging. using CTvox Bruker software.** Whole sample attenuation coefficient can be manipulated in the CTvox software to represent the whole brain in autonomous RGB scales, allowing further contrast to the anatomic features of interest. Therefore, penumbra and core can be highlighted in the 3D context during stroke brain manipulation.

**Suppl. Vid. 2- 3D whole brain of TIA mouse model (tMCAO 10') after osmium tetroxide staining, and micro-CT imaging, using CTvox Bruker software.** Whole sample attenuation coefficient can be manipulated in the CTvox software to represent the whole brain in autonomous RGB scales, allowing further contrast to the anatomic features of interest. Therefore, differences in striatum white fibers between hemispheres can be highlighted (blue) in the 3D context during brain manipulation, with the ipsilateral striatum demonstrating white fiber degeneration when compared to the contralateral side.

**Supplemental Methods:**

**Iohexol staining**

Mice after tMCAO (45min- 24h) were transcardially perfused with 40ml PBS, followed by 40ml of 4% paraformaldehyde (PFA) in PBS. Brains were carefully dissected and post-fixed in 4%PFA, at 4°C, for 1 week. Brains were then rinsed in PBS, and immersed in iohexol (GE Heathcare, Omnipaque 300mg/ml (iodine), 712498.6) diluted 1:2 in PBS (10ml total volume) , according to [1], for 5 days at room temperature, under agitation and protected from light, before micro-CT imaging. As already described, [2], brains were wrapped in parafilm to prevent brain dehydration and scanned using Bruker SkyScan 1276 microCT scanner (Bruker, Belgium). Brain shown in **Supplementary Fig. 1** was acquired with the following settings: 63KV and 200µA, 15µm spatial resolution, 0.4° rotation step through 180°, generating around 514 projections. An Aluminium filter of 0.5mm was used, together with a frame averaging of 4.

**Phosphomolybdic acid staining:**

CD1 Wt mice without any tMCAO surgery were transcardially perfused with PBS and 4% PFA, as previously described, and stained with 2,5% phosphmolybdenic acid (PMA) (Sigma-Aldrich, 221856) diluted in either demineralized 0,9% NaCl (**Suppl. Fig. 4**) or PBS (**Suppl. Fig. 3**) (as described here [3]). Brains were immersed for 7 days, at 4°C, under agitation and protected by light. Brains were prepared for scanning as previously described and scanned with the following settings: 90 KV and 47 µA, 4 µm spatial resolution, 0.2º rotation step through 180 degrees, giving rise to 1801 projections. An Aluminum filter of 1 mm was used, together with a frame averaging of 4. It was clear that PMA only penetrated mouse brain with PBS as solvent. However, the detail of the brain mouse structures is noisier than from the osmium/iodine stain. Even though it performed better than omnipaque and PTA.

**Phosphotungstic acid staining:**

After tMCAO (45min- 24h), mice were transcardially perfused with PBS and PFA, as previously described, and stained with 2.5% phosphotungstic acid (PTA) (Sigma, Ht152) diluted in demineralized water as previously described [4] for 5 days (10mL total volume), at 4°C, under agitation and protected from light. The brain images shown in **Supplementary Fig. 2** were acquired using the following settings: 90KV and 47µA, 8µm spatial resolution, 0.4° rotation step through 180°, originating around 514 projections. An Aluminium filter of 1mm was used, together with a frame averaging of 4.

**Bibliography:**


1. Kuts R, Frank D, Gruenbaum BF, et al. A Novel Method for Assessing Cerebral Edema, Infarcted Zone and Blood-Brain Barrier Breakdown in a Single Post-stroke Rodent Brain. *Frontiers in neuroscience* 2019; 13: 1105-1105. DOI: 10.3389/fnins.2019.01105.
2. Dobrivojevic M, Bohacek I, Erjavec I, et al. Computed microtomography visualization and quantification of mouse ischemic brain lesion by nonionic radio contrast agents. *Croat Med J* 2013; 54: 3-11. 2013/02/28. DOI: 10.3325/cmj.2013.54.3.
3. Dobrivojevic M, Spiranec K, Gorup D, et al. Urodilatin reverses the detrimental influence of bradykinin in acute ischemic stroke. *Exp Neurol* 2016; 284: 1-10. 2016/07/20. DOI: 10.1016/j.expneurol.2016.07.007.
4. Descamps E, Sochacka A, De Kegel B, et al. Soft tissue discrimination with contrast agents using micro-CT scanning. *Belg J Zool* 2014; 144: 20-40.
