## Supplemental figures 1_5 for "High resolution micro-CT imaging in mice stroke models: from 3D detailed infarct characterization to automatic area segmentation"

**Suppl. Fig. 1- Iohexol, phosphotungstic acid (PTA) and phosphomolybdic acid (PMA) staining of whole mice brain.**

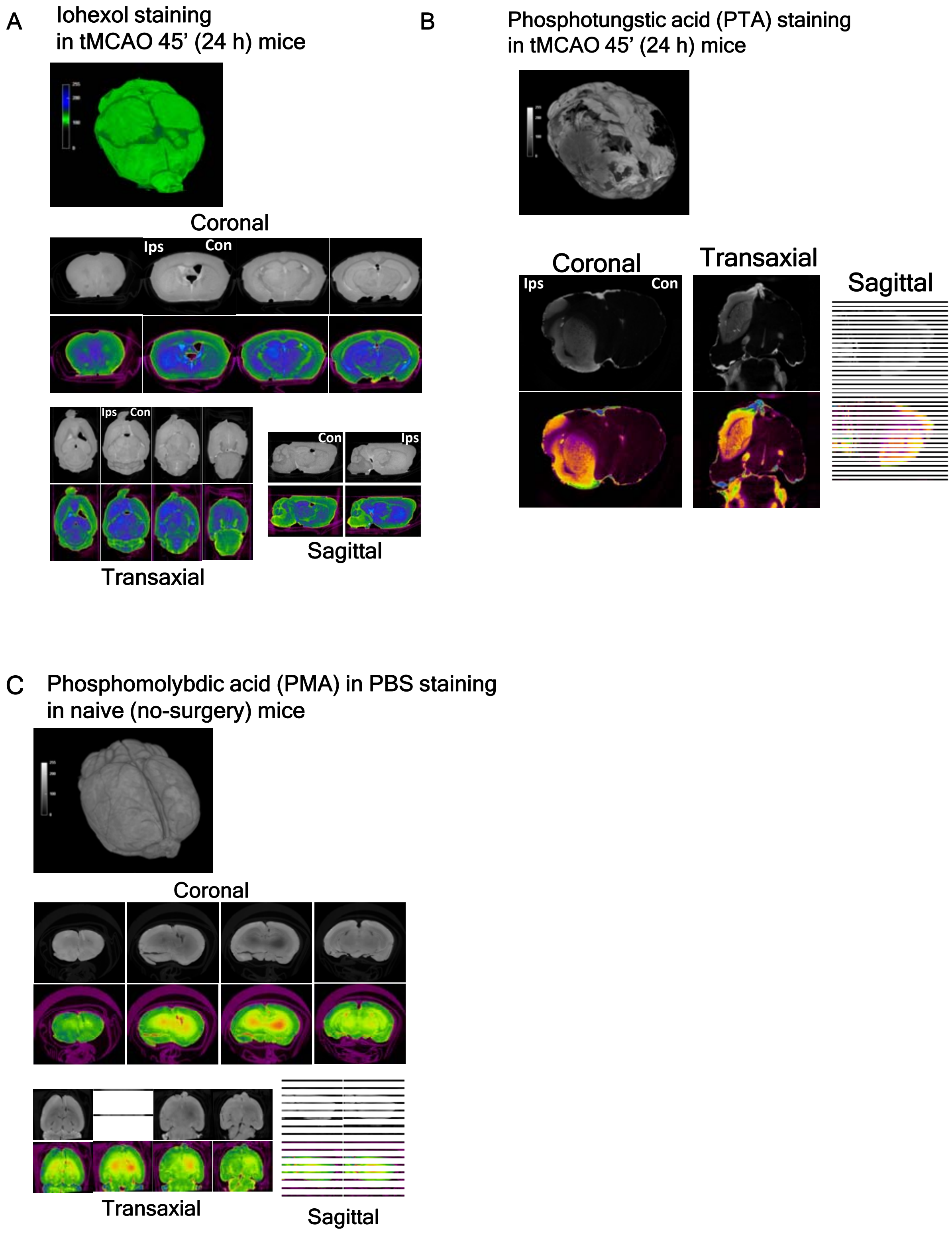

**Suppl. Fig. 2- Iodine staining of mouse brain subjected to allow further immunostaining analysis.**

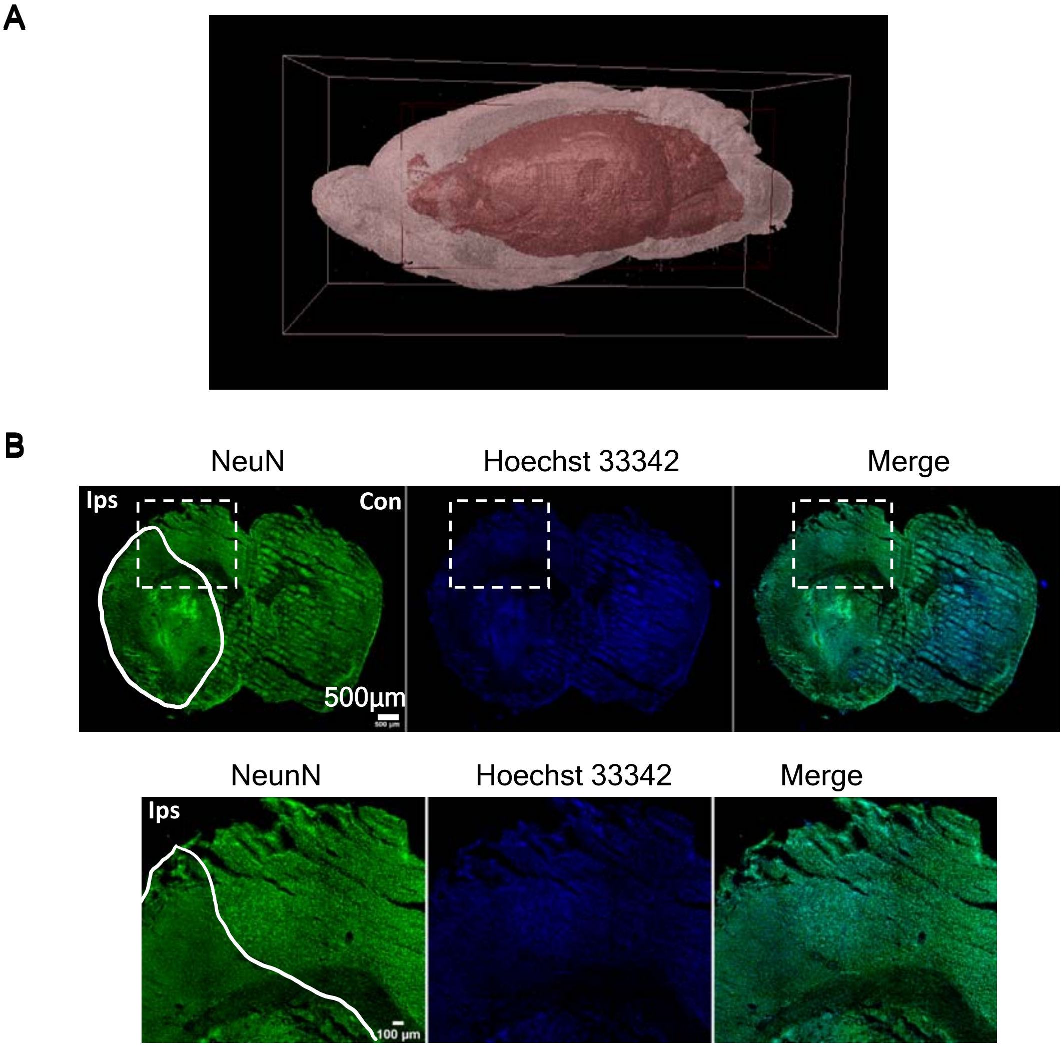

**Suppl. Fig. 3- High resolution micro-CT imaging in mouse stroke model (tMCAO) using iodine staining.**

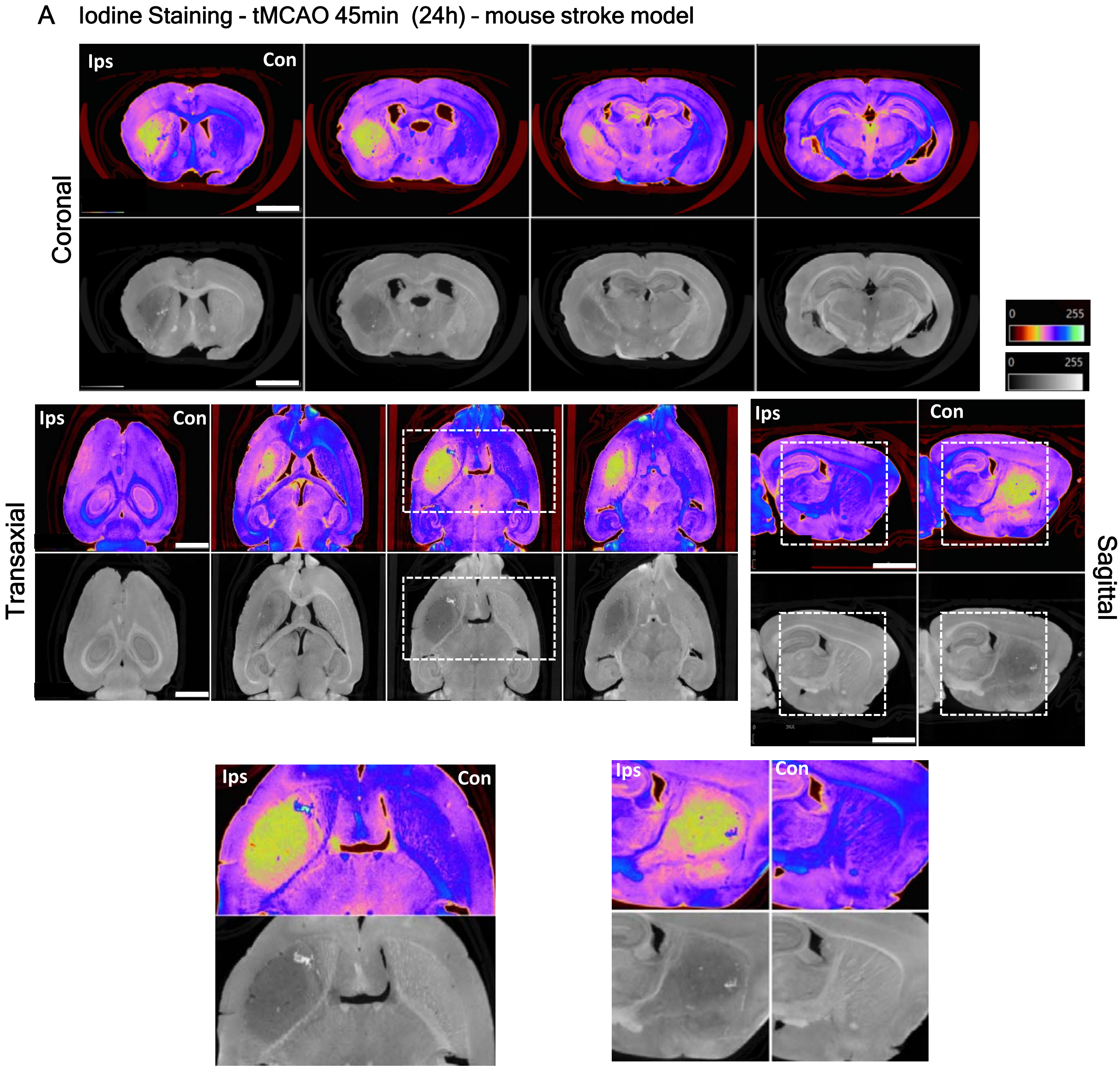

**Suppl. Fig. 4- High resolution micro-CT imaging in TIA mouse model using Osmium Tetroxide staining.**

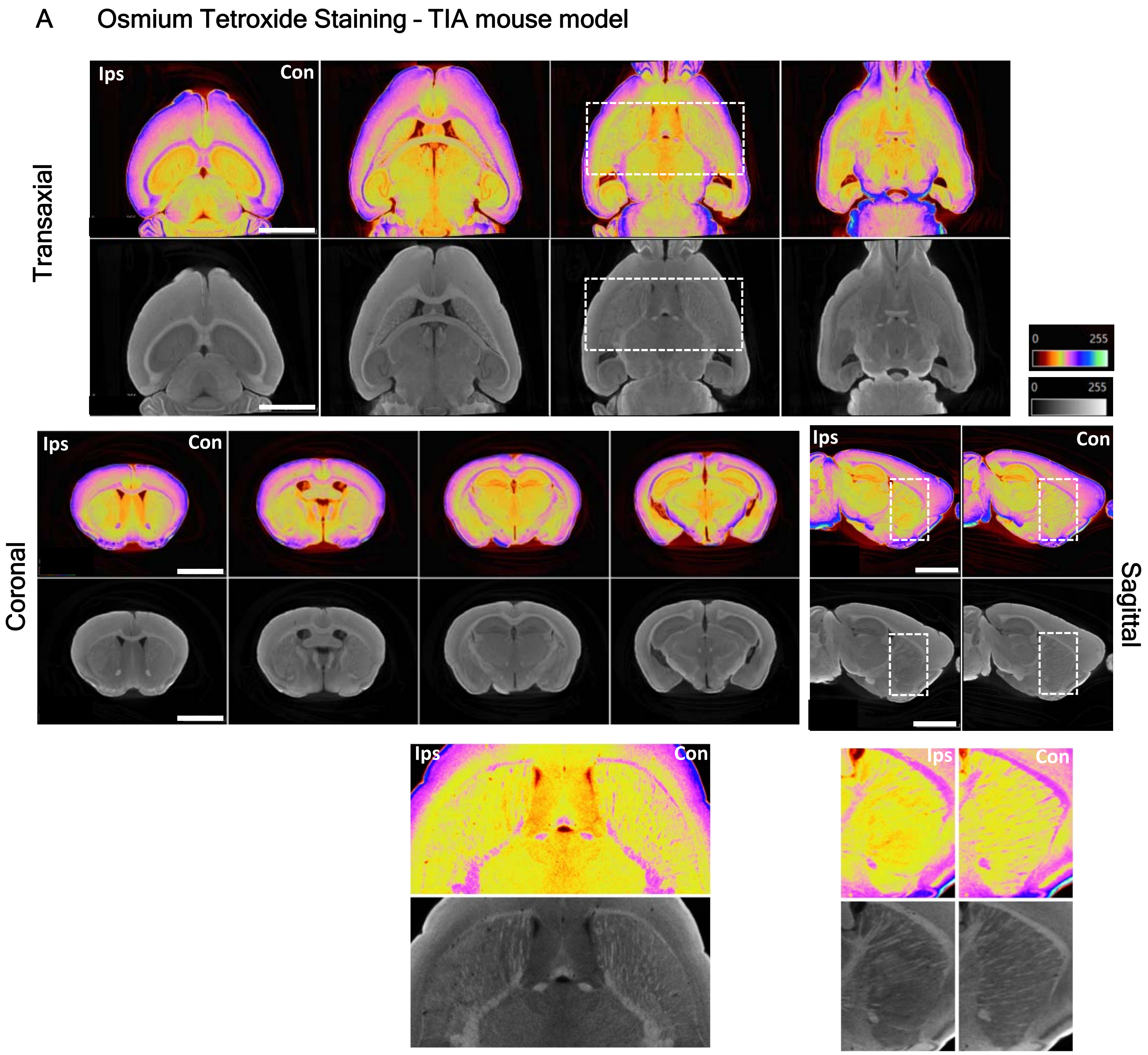

**Suppl. Fig. 5- Brain ischemic lesions (stroke and TIA models) evolution using high resolution micro-CT.**

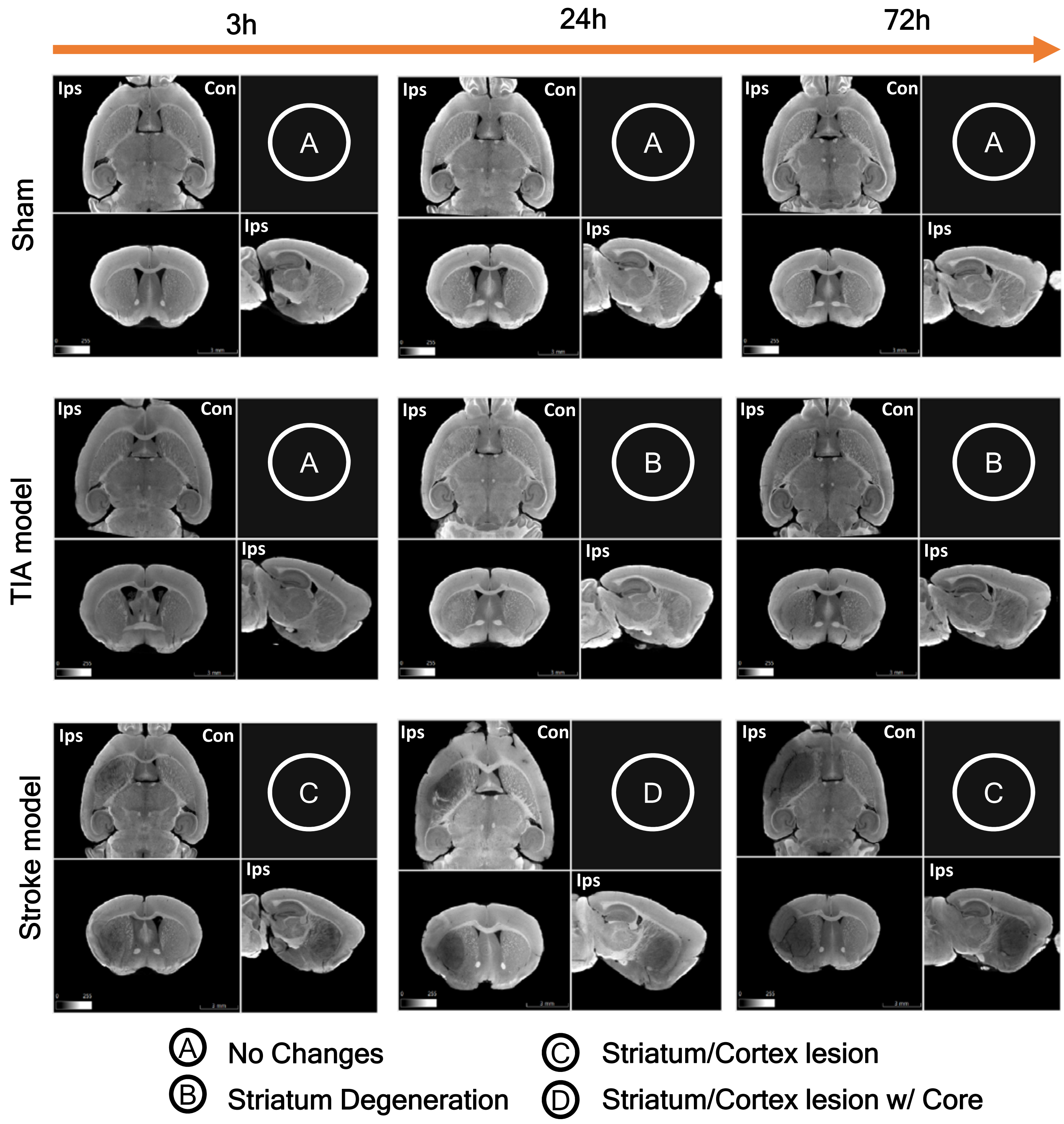
